# A human subcortical connectome at 400 μm resolution

**DOI:** 10.64898/2026.03.31.715699

**Authors:** Chiara Maffei, Ting Gong, Clemens Neudorfer, Dongsuk Sung, Dey Mihir, Kabilar Gunalan, Satrajit Ghosh, Jean C. Augustinack, Susie Y. Huang, Robert M. Richardson, Suzanne N. Haber, Anastasia Yendiki

## Abstract

The fiber pathways of the human subcortex are of particular importance in clinical neuroscience, as they are targeted by neuromodulation therapies, such as deep brain stimulation (DBS) and lesion approaches, in various motor and psychiatric disorders. However, the complexity, size, and position of these pathways make them challenging to image non-invasively with diffusion MRI (dMRI). As a result, recent efforts in atlasing these pathways to guide neuromodulation have resorted to using synthetic data. Here we present the first extensive reconstruction of fiber pathways of the human subcortex with ultra-high-resolution dMRI. We leverage a unique, *ex vivo* dMRI dataset acquired by the BRAIN CONNECTS center for Large-scale Imaging of Neural Circuits (LINC) on the first-of-its-kind Connectome 2.0 scanner. Our contribution is two-fold. First, by showing the feasibility of reconstructing these pathways with non-invasive neuroimaging at the single subject level, we set the stage for future research into reconstructing them *in vivo* in individual patients. Second, we provide a high-definition atlas of basal-ganglia-thalamocortical circuits that is readily extensible via our publicly released data and annotations. As a first demonstration of its clinical validity, we align our atlas to “hotspots” from previous DBS studies and identify pathways associated with therapeutic or side effects.

## 1. Introduction

Localizing the complex pathways of the human subcortex noninvasively is critical for targeting them in neuromodulation therapies for various motor and psychiatric conditions, including Parkinson’s Disease (PD), dystonia (DYT), essential tremor (ET), obsessive compulsive disorder (OCD), and Tourette’s Syndrome (TS). Deep brain stimulation (DBS) in particular, which modulates primarily myelinated axons and thus white matter tracts^1–3^, is a key clinical application where advancing our ability to image these tracts promises not only to improve targeting^4–6^, but also to provide post-hoc insights into the networks mediating therapeutic benefit^7–10^. However, the resolution and signal-to-noise ratio (SNR) of conventional diffusion MRI (dMRI) has been insufficient for a detailed reconstruction of the fiber pathways implicated in these therapeutic interventions. As a result, classical histological atlases remain the principal reference on the complex trajectories of these pathways and their relationship with surrounding grey matter nuclei.

Common neurological and psychiatric DBS targets focus on the connections of basal ganglia (BG) structures: the striatum, globus pallidus (GP), substantia nigra (SN) and subthalamic nucleus (STN). These connections include those of the GP (for PD and DYT), the STN (for PD and increasingly for OCD), and the thalamic nuclei (for ET and TS). The complex connections of the BG are often summarized by three main circuits: the direct, indirect, and hyperdirect pathways (Fig. 1). The direct pathway connects the striatum with the internal segment of the GP (GPi) and the SN, which, in turn, project to the thalamus and back to cortex. The indirect pathway connects the striatum with the external segment of the GP (GPe), which projects to the STN, and which, in turn, projects to the GPi. The hyperdirect pathway relays cortical information directly to the STN^11^ (Fig. 1).

**Figure 1.**
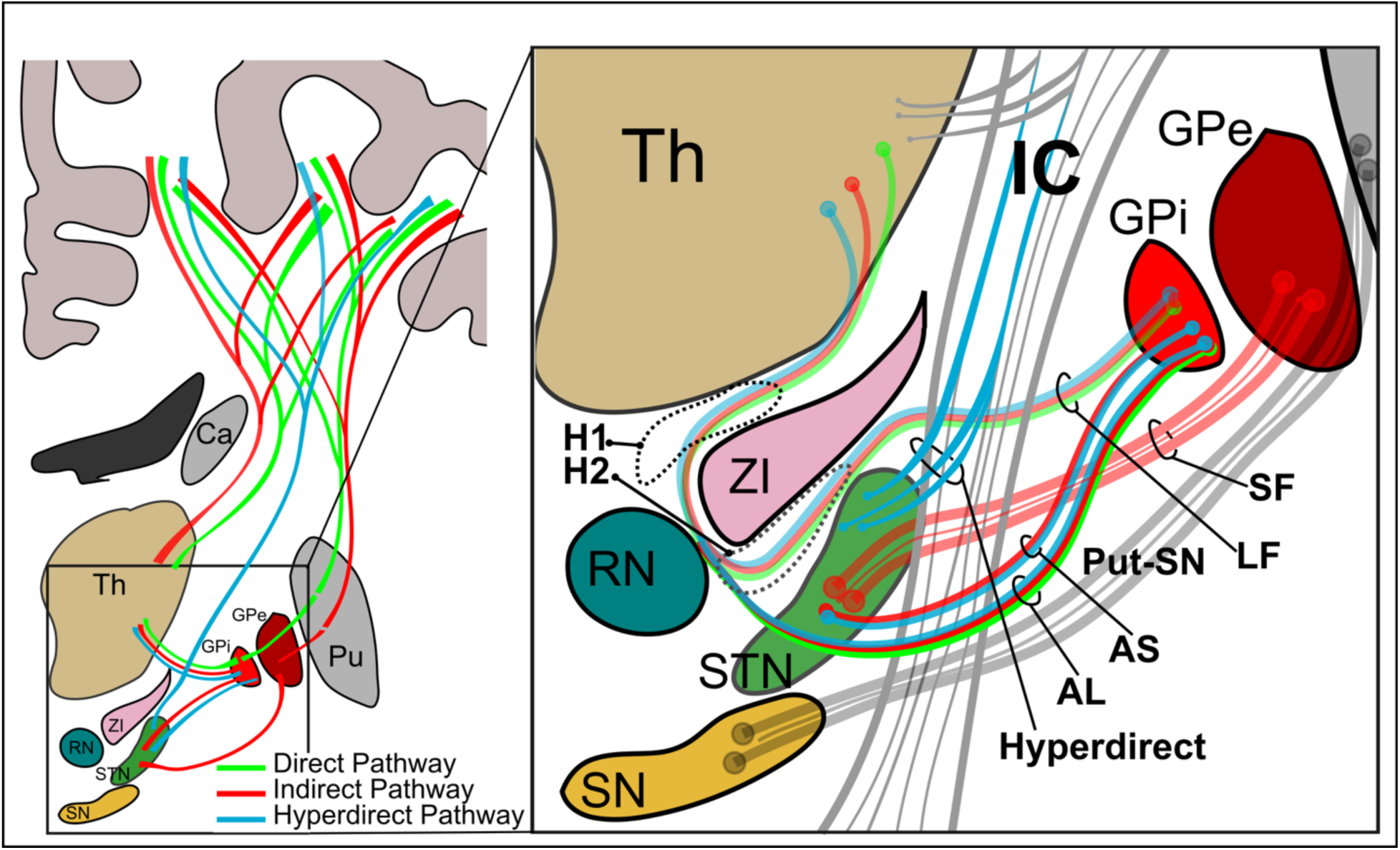
Schematic of basal ganglia pathways. **Left:** Schematic of the direct (green), indirect (red), and hyperdirect (blue) basal ganglia loops. The direct pathway includes connections from the cortex to the striatum (here putamen (Pu)), to the globus pallidus internus (GPi), to the thalamus (Th), and then back to the cortex. The indirect pathway includes connections from the cortex to the striatum (here putamen), to the external segment of the globus pallidus (GPe), to the subthalamic nucleus (STN), back to the GPi, thalamus and back to the cortex. The hyperdirect pathway sends cortical inputs straight to the STN, and from here to the GPi and thalamus and back to the cortex. The zona incerta (ZI), red nucleus (RN), caudate (Ca), and substantia nigra (SN) are also shown. **Right:** Blow up of the pallidal pathways projecting to the thalamus and STN and the pathways they intersect. AL: ansa lenticularis; AS: ansa subthalamica; GP: globus pallidus; H1/H2: Forel’s fields; IC: internal capsule; LF: lenticular fasciculus; Put: Putamen; RN: red nucleus; SF: subthalamic fasciculus; SN: substantia nigra; STN: subthalamic nucleus; Th: thalamus; ZI: zona incerta.

Invasive anatomical studies in both humans and non-human primates have described the connectivity of these intermingling fiber pathways in detail^12–17^. GPi-thalamic axons travel in the ansa lenticularis (AL) and lenticular fasciculus (LF). GPe axons project to STN via the subthalamic fasciculus (SF), and STN-GPi axons travel within the ansa subthalamica (AS). Together, these pathways form a dense network of crossing and overlapping fibers within the subthalamic region (inset in Fig. 1), where pallidal projections course through and around the internal capsule (IC), while swirling around the STN, ZI, and RN, and intersecting with fibers of the hyperdirect and the striatonigral (Put-SN) pathway. This anatomical complexity, combined with the deep location and small size of these fiber bundles, make them very challenging to resolve with conventional noninvasive neuroimaging. Reflecting this limitation, recent efforts to build atlases of BG pathways relevant to neuromodulation have resorted to generating synthetic, idealized models of these fiber bundles^18,19^, rather than reconstructing them from real neuroimaging data.

The present work overcomes these limitations by leveraging the highest-resolution dMRI data acquired in whole *ex vivo* human hemispheres to date. These data, collected as part of the BRAIN CONNECTS center for Large-scale Imaging of Neural Circuits (LINC) on the first-of-its-kind Connectome 2.0 scanner, allow us to perform the first comprehensive reconstruction of human BG pathways with dMRI tractography at the single subject level. We map these reconstructions from the first two specimens to a common coordinate framework, and create the most comprehensive, dMRI-based atlas of human subcortical pathways to date. The atlas and imaging data are available publicly through https://gallery.lincbrain.org, and will be updated as additional specimens are imaged.

In the following, we present three analyses to demonstrate the validity of our atlas. First, we map the topographic organization of BG and thalamo-cortical projections and replicate, for the first time in the human brain, gradients previously reported in non-human primate studies. Second, we compare BG pathways and additional surrounding brainstem and limbic pathways, as reconstructed from our ultra-high-resolution *ex vivo* dMRI, to those reconstructed using conventional *in vivo* dMRI data from the Human Connectome Project (HCP), demonstrating a significant methodological advance over the state-of-the-art. Third, as an illustration of the utility of our tract atlas for connectomic neuromodulation, we use it to analyze data from previous DBS studies and identify pathways associated with therapeutic and side effects of DBS. Connectomic fingerprints from this analysis demonstrate a domain-specific correspondence between patterns of clinical improvement and associated circuits.

Reconstructing these subcortical circuits with high-resolution *ex vivo* dMRI tractography represents a first, critical step toward *in vivo* mapping in individual patients. In addition to the clinical utility of our atlas as a normative connectome for both prospective targeting and retrospective analysis of neuromodulation in a common coordinate framework, we expect the high-quality data and pathway annotations from our *ex vivo* dMRI acquisitions to fuel future research into training and validating models for reconstructing these pathways from *in vivo* dMRI.

## Results

### Extensive tractography-based reconstruction of the classical basal ganglia circuits

The classical direct, indirect, and hyperdirect circuits of the BG are typically depicted in schematic diagrams like the ones in Fig. 1. The unprecedented spatial resolution of the LINC dMRI data allows us to delineate both short- and long-range connections within these three circuits in their entirety, and hence to produce their first complete, 3D reconstruction in the human brain (Fig. 2). The long-range thalamo-cortical and striato-cortical connections are easier to identify with tractography given their larger size and have thus been reconstructed before (Fig 3a). In contrast, the shorter-range components interconnecting the STN, GP, and thalamus have not been comprehensively reconstructed in the human brain before. These connections are composed of pallidofugal fibers projecting to the STN, thalamus, habenula (Hb), and midbrain regions. Here, we tease apart all these different components, including the classical ansa subdivisions (AL, LF, SF) and the more recently described ansa subthalamica (AS) (Fig. 3b-c). This level of detail is critical for accurately modeling the effects of neuromodulation on these circuits.

**Figure 2.**
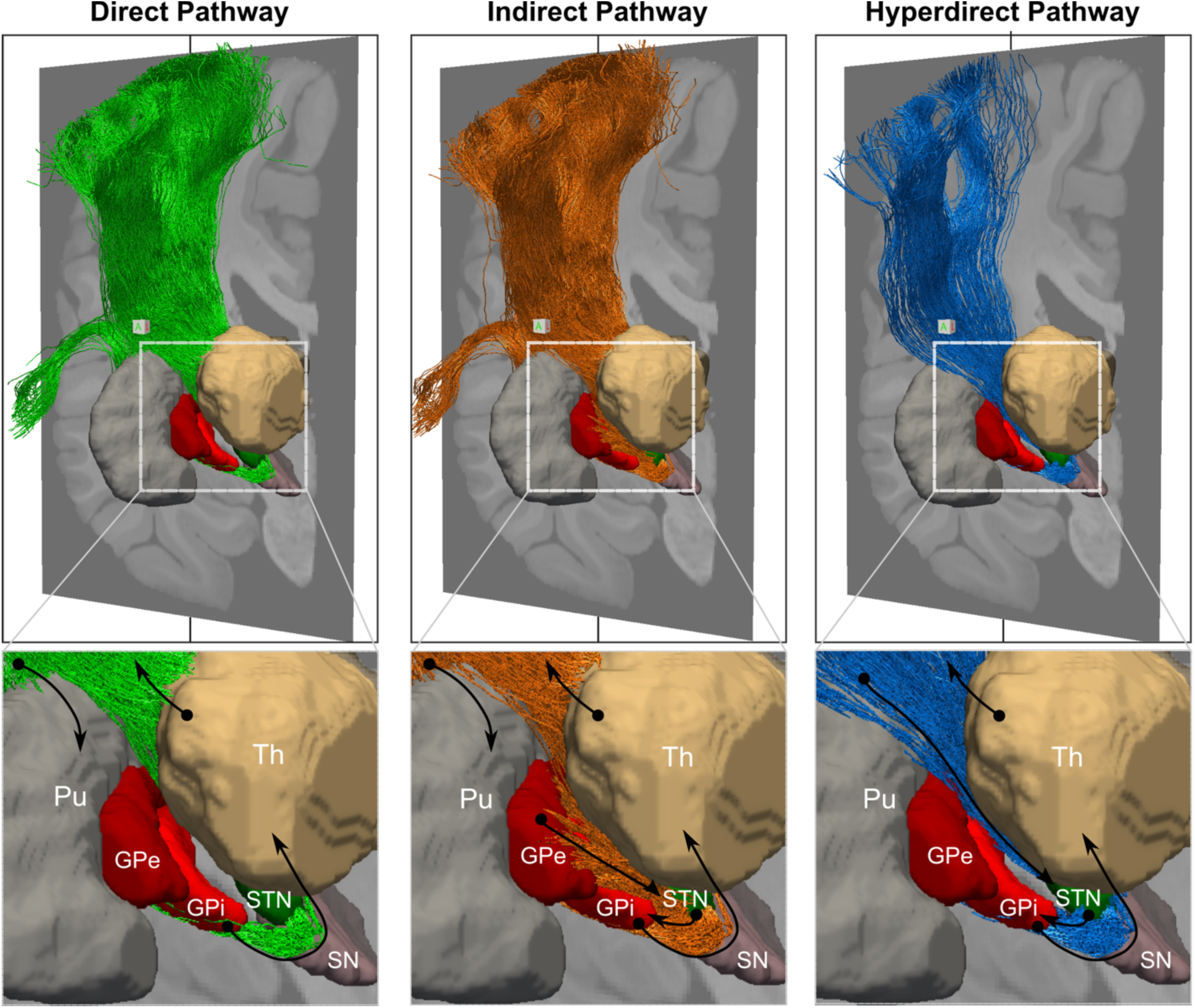
Tractography-based reconstruction of the basal ganglia circuits. The figure shows the reconstruction of the direct (green), indirect (orange), and hyperdirect (blue) pathways of the basal ganglia. Only the motor (cortical area 4) and premotor (cortical area 6) components of the striato-cortical, thalamo-cortical, and hyperdirect projections are shown. Close ups focus on the subcortical components of the pathways with black arrows indicating the flow of information within the circuit, from the cortex to the putamen, through the basal ganglia, to the thalamus, and back to the cortex. The SN is also shown in sand. GP: globus pallidus; Pu: Putamen; SN: substantia nigra; STN: subthalamic nucleus; Th: thalamus.

**Figure 3.**
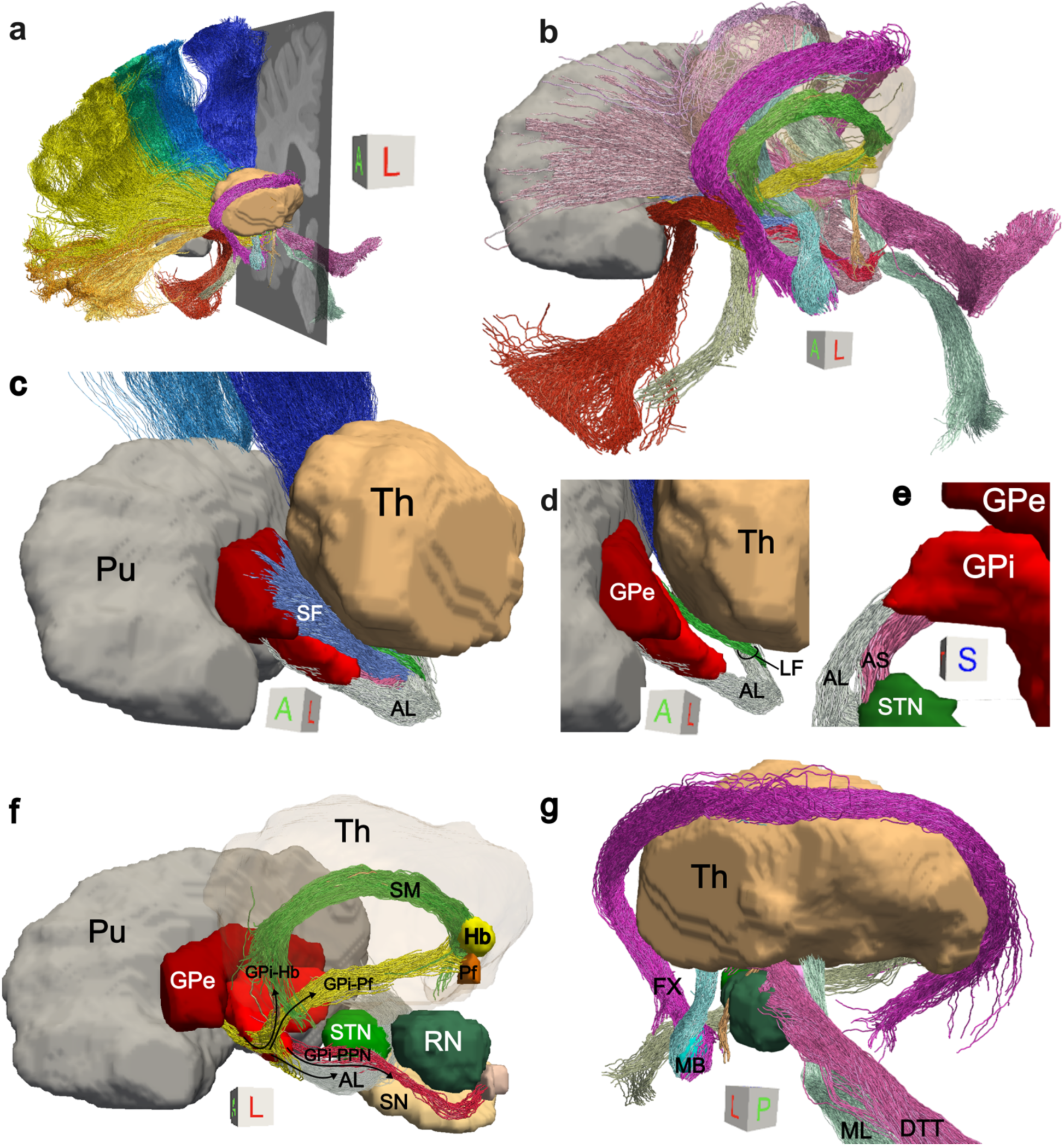
Tractography atlas of subcortical pathways. **a)** Overview of the atlas. 3D reconstructions for all pathways are shown together with 3D renderings of putamen (grey) and thalamus (sand). **b)** 3D reconstructions for subcortical pathways only. **c)** Rendering of BG fiber systems, illustrating the separation between the AS (pink), AL (grey), LF (green) and SF (blue), together with their spatial relationships with the thalamus and GP. **d)** Close-up of AL and LF pathways as they leave the GPi and enter the thalamus. **e)** Superior view of the AS and AL as they leave the GPi. **f)** GPi projections to the Pf, the Hb, and the PPN. **g)** Posterolateral view of pathways surrounding core BG circuits. Caudally, the DTT and ML carry information from the cerebellum and brainstem, respectively. More anteriorly, the MTT and FX ascend from the MB and the FR runs medial to the RN. AL: ansa lenticularis; AS: ansa subthalamica; DTT: dentatothalamic tract; FX: fornix; GP: globus pallidus; Hb: habenula; LF: lenticular fasciculus; MB: mammillary bodies; ML: medial lemniscus; MTT: mammillothalamic tract; Pf: parafascicular nucleus of the thalamus; Pu: Putamen; RN: red nucleus; SF: subthalamic fasciculus; SN: substantia nigra; STN: subthalamic nucleus; Th: thalamus; ZI: zona incerta. A comprehensive list of abbreviations can be found in the supplementary materials.

Our dMRI-based reconstructions reflect descriptions of these circuits from prior, invasive anatomic studies. The AL arises from the rostral, outer GPi and courses ventromedially to the medial GPi and anterior to the IC, while the LF arises from the caudal, medial GPi and crosses the IC before reconnecting with the AL (Fig 3d). The AS runs lateral to the AL and connects anteroventral GPi to anteroventral STN (Fig 3e). In addition to these classical subdivisions, we reconstruct pallidal connections to the Hb and parafascicular nucleus of the thalamus (Pf) (GPi-Pf, GPi-Hb) that heretofore have only been described in classical anatomic and tracer studies^16,20^. The GPi-Hb fibers separate from the main body of the ansa near the apex of the GP, climb dorsally over the rostral curvature of the thalamus and join the stria medullaris (SM)^14^ (Fig. 3f). GPi-Pf fibers leave the GPi and traverse the thalamus mediodorsally to terminate in the thalamus Pf^16^, located infero-lateral to the habenula. The caudal-most target of pallidal fibers is the pedunculo-pontine nucleus (PPN)^14^, located inferior, posterior lateral to red nucleus (RN) and dorsal to the substantia nigra (SN)^17^. The GPi-PPN pathway is formed by fibers that separate from Forel’s field H2 and collect dorsal and medial to the STN^14^ before descending towards the pons (Fig. 3f).

### A high-precision atlas of pathways of the human subcortex

In addition to the core BG pathways described above, our ultra-high-resolution dMRI data allow us to reconstruct several challenging limbic and brainstem pathways that course around the BG. The medial lemniscus (ML) and dentato-thalamic tract (DTT) enter the tegmental region caudally, run through the subthalamic area and project to the ventral posterior and lateral thalamus, respectively. Medially, the fasciculus retroflexus (FR) runs within the medial aspect of the RN to converge with the SM in the Hb, and more anteriorly the mammillo-thalamic tract (MTT) and fornix ascend from the mammillary bodies (MB) (Fig. 3g). While most of these pathways have diameters smaller than 2mm^21^, we provide accurate reconstructions using high-resolution tractography. In total, we provide annotations of 44 pathways within the basal forebrain, mesencephalon, and diencephalon, along with 10 manually labeled subcortical nuclei (Supplementary Fig. 1). While results are shown here in individual space, we also provide all nuclei and tracts mapped to the MNI-2009b-ICBM template, to facilitate their use in neuroimaging and neuromodulation studies.

### Tractography replicates topographic gradients known from invasive studies in non-human primates

The BG pathways are generally topographically organized, such that motor, cognitive, and limbic systems project to distinct subzones within each structure, including the STN^22^. The high resolution and SNR of our dMRI data allow us to disentangle the projections from the motor, premotor, prefrontal, and orbitofrontal cortex and follow their trajectory as they travel through the IC (Fig. 4a). The organization of streamlines within the IC replicates topographic rules from invasive tracer studies, with dorsal prefrontal and premotor streamlines running superior to ventral prefrontal and orbitofrontal fibers^23–25^. Of note, we successfully map small projections from cortical regions that are challenging to reconstruct from conventional dMRI data (*e.g*., Opro and area 10v) (Fig. 4a, Supplementary Fig. 2). In agreement with tracer studies in macaques^26^, caudal to the anterior commissure (AC), streamlines from each cortical region split in two components: one that enters the thalamus, and one that continues down to the STN and brainstem. We replicate these results in a second brain specimen (Supplementary Fig. 3).

**Figure 4.**
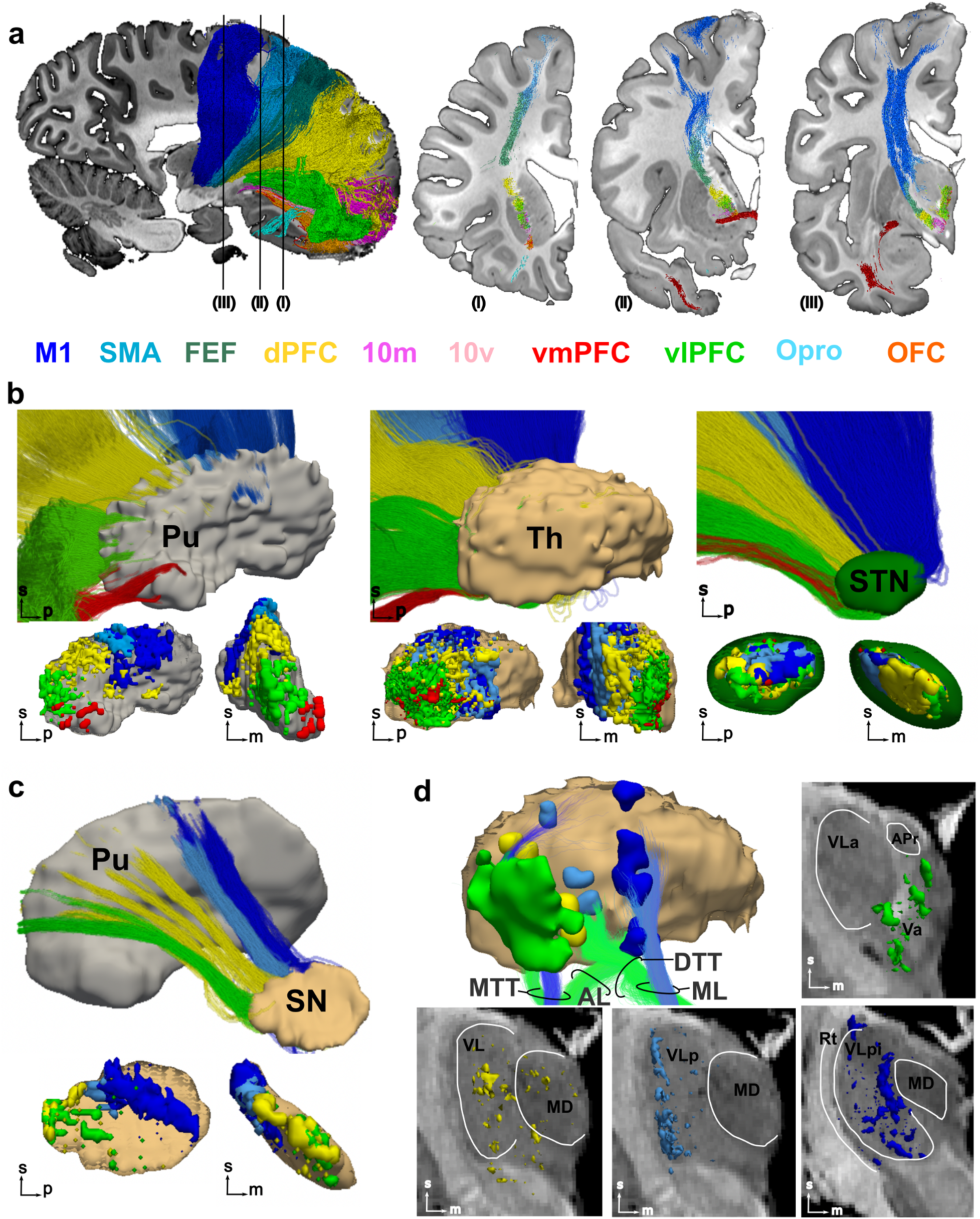
Topographic organization of BG pathways. **a)** Segmentation of motor, premotor, prefrontal, and frontal fibers running through the internal capsule (IC). 3D reconstruction of the different components (left) together with 2D coronal sections showing the location of the different components within the IC and their relationship to the anterior commissure (in dark red). The approximate location of the coronal slices is indicated by roman numbers. Streamlines are colored based on the cortical region they project to. Results are shown for one sample. Supplementary Fig. 3 shows results for a second sample. **b)** Top: Sagittal views of cortico-striatal (left), cortico-thalamic (middle), and cortico-subthalamic streamlines segmented based on the cortical region they project to. Bottom: close-ups of streamline termination distributions within the putamen (left, in grey), the thalamus (middle, in light brown), and STN (right, in green) in both sagittal and coronal planes (orientation indicated by arrows in the bottom left corner of each panel). Terminations are color-coded based on cortical regions. **c)** Segmentation of striato-nigral streamlines generated using cortico-striatal terminations (top). The streamline terminations obtained from these segmentations are mapped within the SN and shown in sagittal and coronal views (bottom). **d)** The largest clusters of cortical terminations within the thalamus are shown together with major thalamic afferent pathways (top left). The full distribution of cortical terminations for motor, premotor, dorsal and ventral PFC are shown on coronal slices highlighting their accurate localization within motor and associative thalamic nuclei, respectively. dlPFC: dorsolateral PFC; dmPFC: dorsomedial PFC; DTT: dentatothalamic tract; MD: medial dorsal nucleus of thalamus; ML: medial lemniscus; MTT: mammillothalamic tract; STN: subthalamic nucleus; Th: thalamus; VL: ventrolateral nucleus of thalamus; vlPFC: ventrolateral PFC; vmPFC: ventromedial PFC.

For the first time with dMRI, we show the feasibility of precisely mapping the terminations of the cortico-striatal, cortico-thalamic, and cortico-subthalamic pathways within each of their subcortical targets (Fig. 4b). In the putamen, we observe a ventromedial to dorsolateral gradient: prefrontal projections are concentrated anterior to the AC primarily within the ventral and rostral putamen, while motor and premotor regions project to dorsolateral and central sections caudal to the AC^27,28^. Streamlines from vmPFC are sparse and confined to the rostral ventromedial putamen, while vlPFC streamlines terminate along the ventral rostral putamen^29^. The thalamus exhibits a rostrocaudal gradient, with PFC terminations located rostral to premotor and motor terminations. The STN exhibits a rostrocaudal and ventromedial to dorsolateral organization, with dPFC terminations concentrated in the medial half of the nucleus, whereas motor terminations concentrate dorsally in the lateral half, and premotor medial to motor^22^. In agreement with tracer studies in non-human primates, we identify only sparse STN-vmPFC streamlines, correctly located within the medial tip of the STN^22^. We further investigate the topographic organization of the BG circuits by using cortico-striatal terminations within the putamen to segment striato-nigral projections and map topography within the SN (Fig. 4c). Striatal terminations within the SN reflect the ventrolateral dorsomedial gradient described in tracer studies^30^, with motor and premotor terminations concentrated within the ventral lateral caudal portions of the SN and frontal regions distributed more mediodorsally.

While the distribution of axonal terminations within the putamen, STN, and SN show partial overlap across functional domains, output thalamic afferents relay to distinct thalamic nuclei associated with motor, associative, and limbic loops^31^. We investigate this by assessing the relationship between the largest clusters of cortico-thalamic terminations within the thalamus and key thalamic afferents, including the MTT, the AL, the ML, and the DTT (Fig. 4d). The ML, which primarily targets the ventral posterior lateral (VPL) motor thalamic nucleus, overlaps with motor cortico-thalamic terminations. The MTT, whose main thalamic target is the ventral anterior nucleus (VA), overlaps with vlPFC terminations, which distribute over the VA and medial dorsal thalamic nucleus (MD). Similarly, AL and DTT terminations are localized within the VL and overlap with premotor and prefrontal clusters, demonstrating the precise topography of thalamic afferents relative to cortical projections.

### Anatomic fidelity of tractography in our ex vivo dMRI data vs. HCP data

Our tractography reconstructions represent a significant advance in anatomic accuracy over the state-of-the-art public dMRI data from the HCP^32^, which are often leveraged in connectomic studies. We highlight this by comparing a subset of the labeled pathways, chosen to include both short- and long-range pathways of different size and anatomical complexity. Figure 5 shows that several of the small, challenging pathways (FR, AL, AS, MTT) cannot be reconstructed at all using the HCP data (Fig. 5a-b). At this resolution (1.55 mm) the signal is dominated by the large thalamic and cerebellar pathways projecting to the frontal cortex (Fig 5c-d). Other larger pathways (DTT, ML, STN-4, STN-6, FX), while reconstructed, show lower anatomical accuracy compared to the LINC data. For example, the DTT and ML show sparser streamlines with less accurate thalamic terminations (Fig. 5e) and the premotor component of the hyperdirect pathway (STN-6) show less streamlines correctly exiting the entering the STN (Fig 5f).

**Figure 5.**
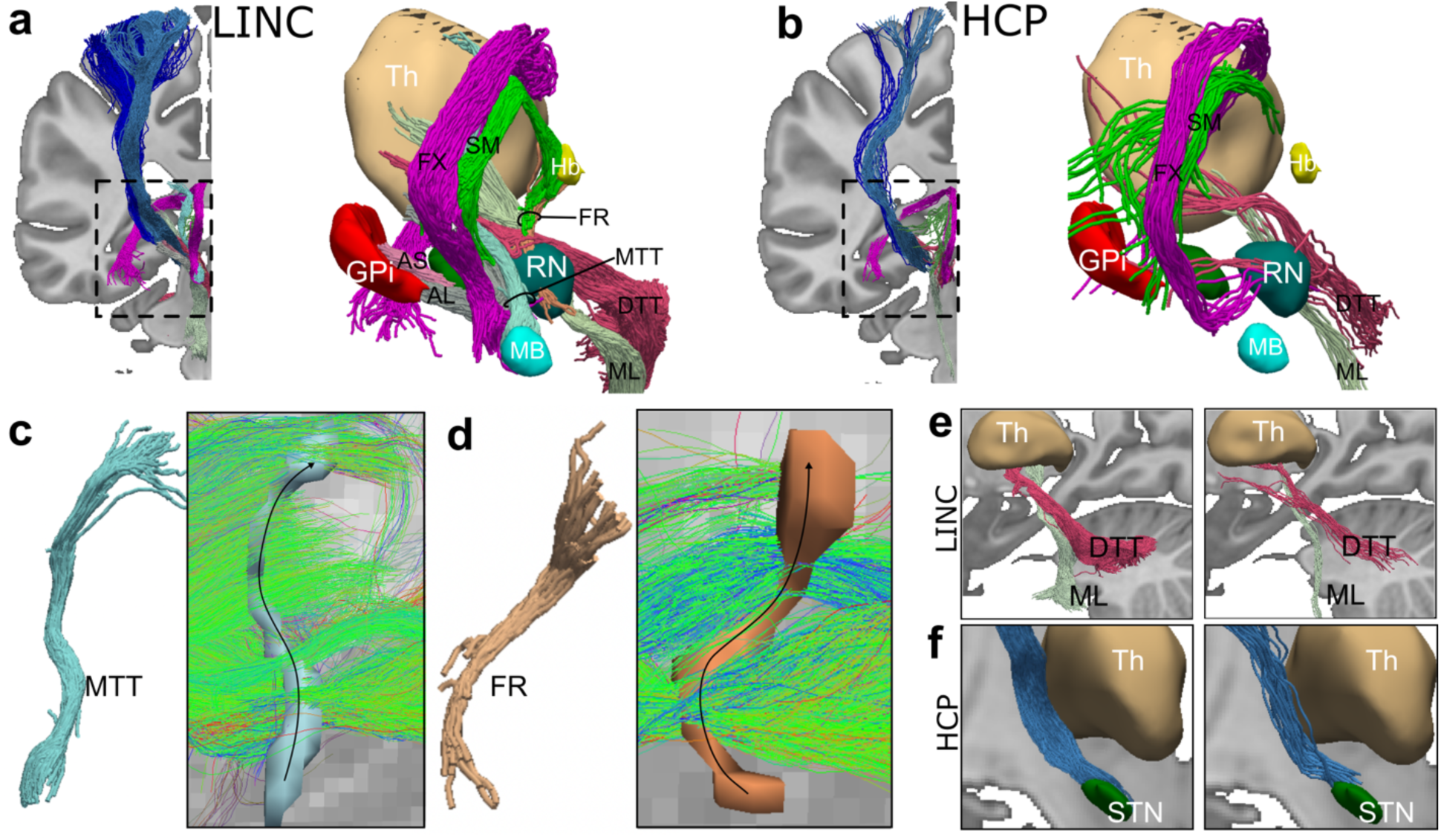
Comparison of LINC to HCP data. Tractography reconstructions generated with LINC **(a)** vs HCP **(b)** data. For each dataset, coronal views of tractography reconstructions are shown in MNI-2009b-ICBM space overlaid on the T1-weighhted volume. A close up of the pathways running within the subthalamic region shows several short tracts are missing from the HCP dataset **c-d)** Set of streamlines generated from the HCP data when using the visitation maps of the MTT and FR extracted from the LINC atlas as inclusion ROIs. Streamlines are color coded based on orientation (green: anterior-posterior; red: right-left; blue: inferior-superior). Black arrows superimposed on visitation maps indicate the correct direction of fibers within the FR and MTT. These images show that the HCP data are dominated by streamlines oriented anterior-posterior and do not recover fibers within the FR and MTT. **e)** Close up of the DTT and ML reconstruction for the LINC vs HCP data showing improved accuracy of their terminations within the thalamus for the LINC data**. f)** Close up of the premotor component of the hyperdirect pathway showing improved accuracy of STN terminations for the LINC data. AL: ansa lenticularis; AS: ansa subthalamica; DTT: dentatothalamic tract; FX: fornix; GP: globus pallidus; Hb: habenula; MB: mammillary bodies; ML: medial lemniscus; MTT: mammillothalamic tract; RN: red nucleus; STN: subthalamic nucleus; Th: thalamus. A comprehensive list of abbreviations can be found in the supplementary materials.

### Connectomic fingerprints of therapeutic effects and side effects in DBS

In a first demonstration of the utility of our *ex vivo* tract atlas as a normative connectome for analyzing *in vivo* DBS studies, we generate connectivity fingerprints associated with both clinical improvement and stimulation-induced side effects. We use “hotspots” from the DBS literature (Supplementary Table 1), aligned to MNI-2009b-ICBM template space, and compute their overlap with the pathways in our atlas. Each hotspot overlaps with a distinct set of pathways, with varying degrees of involvement across motor, psychiatric, and autonomic domains (Fig. 6 and Supplementary Fig. 5).

**Figure 6.**
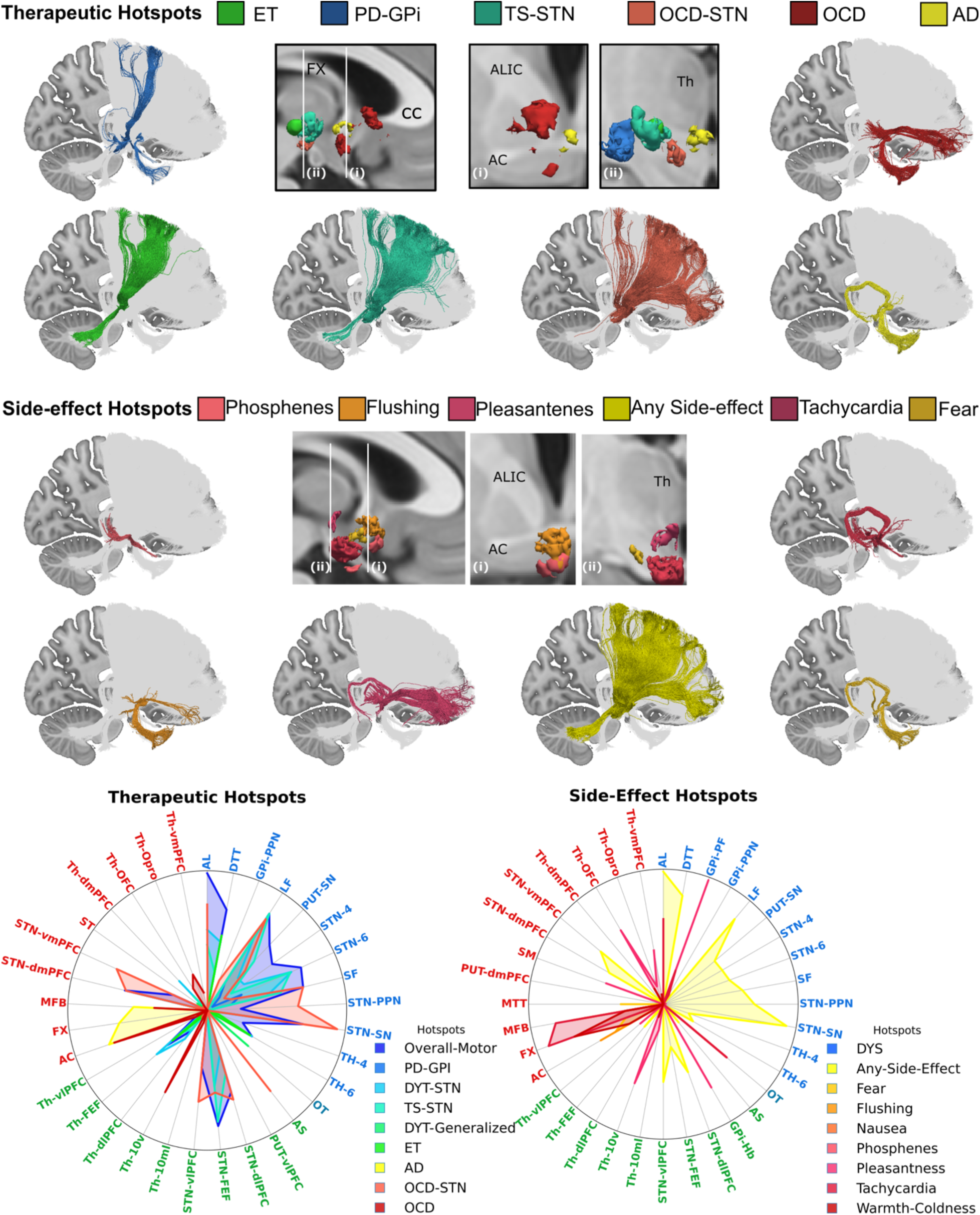
Connectivity fingerprints of therapeutic and stimulation-induced side effects. For both therapeutic and side-effect hotspots, 3D volumes from a subsample of the hotspot maps are shown in the middle in sagittal view and at two different coronal levels (more posterior (i) and more anterior (ii)). Tracts overlapping with each volume are displayed color-coded based on associated hotspot, whereas the non-overlapping tracts in the atlas are shown in grey. Radial plots show the fraction of involvement of each tract for all therapeutic and side-effect hotspots. Tract labels are colored based on whether they belong to the motor (blue), sensory (light blue), limbic (red), or associative (green) systems. AD: Alzheimer’s disease; DYT: dystonia; ET: ET: essential tremor; GPi: internal segment of the globus pallidus; OCD: Obsessive Compulsive Disorder; PD: Parkinson’s Disease; STN: subthalamic nucleus; TS: Tourette’s Syndrome.

Overall, hotspots associated with reduction in motor symptoms show the most widespread engagement across motor pathways, while non-motor symptoms exhibit more focal and selective involvement of sensory, associative, and limbic pathways (Supplementary Fig. 6). Hotspots associated with overall improvement of motor symptoms (both rigidity and akinesia) in PD^33^ intersect broadly with BG pathways, with prominent engagement both its cortical (STN-4, STN-6, STN-FEF) and subcortical components (STN-PPN, SF, AL, LF, STN-SN). Essential tremor improvement ^34^ is characterized by a more focal network architecture that primarily involves the DTT and thalamic connections to premotor regions, consistent with previous reports^35^ (Fig. 6). Stimulation of the GPi for PD^36^ primarily engages cortical and subcortical components of the BG pathways (LF, SF, STN-6), but also shows some overlap with the AC and OT, pathways highly involved in side-effect networks (Fig. 6). Hotspots linked to clinical improvement in OCD involve different networks depending on the stimulation target. Stimulation of the ventral IC^29^ overlaps with the AC, MFB and the limbic components of the thalamo-cortcial projections (Th-10v, Th-Opro, Th-vmPFC). In contrast, STN targeting engages several cortical and subcortical BG pathways, including striato-nigral pathways, AL, SF, STN-PPN, and AS. Although there is overlap with STN stimulation sites that improve motor symptoms^27^, the STN hotspot for OCD^10^ shows greater involvement of dorsal and ventral PFC connections and selective involvement of GPi-PPN and AS (Fig. 6). Finally, networks associated with therapeutic effects in Alzheimer’s disease are primarily centered on the FX, a key memory pathway within the Papez circuit, but also involve the AC and, to a lesser extent, the MFB.

Autonomic side-effects (SE) ^37^ are associated with selective involvement of hypothalamic and limbic pathways. Nausea and phosphenes demonstrate highly focal involvement of the OT, consistent with sensory activation. Fear-related and warmth-cold side effects are both associated with involvement of the FX, AC, and MFB, but fear also shows moderate involvement of the MTT. Flushing involves the AC in conjunction with the MFB and vmPFC, suggesting engagement of limbic–autonomic circuitry. Pleasantness shows strong involvement of the vmPFC and habenula connections, including the SM and GPi-Hb. Compared to the hotspot associated with overall improvement of motor symptoms in PD ^33^, the related hotspot eliciting any side effect in PD demonstrates relatively greater involvement of both dorsal and ventral PFC connections, indicating a shift toward associative and limbic network engagement.

Together, these findings reveal distinct connectomic fingerprints underlying therapeutic benefit and stimulation-induced side effects. Motor improvement is associated with widespread engagement of basal ganglia–thalamo–cortical motor networks, whereas psychiatric and autonomic effects arise from more selective involvement of limbic, associative, and hypothalamic pathways. The consistency and anatomical validity of these pathway–symptom relationships across diverse clinical phenotypes underscore the anatomical accuracy of the atlas and demonstrate its utility for identifying specific white matter tracts that mediate both efficacy and adverse effects in DBS.

## Discussion

This work makes two main contributions. First, we demonstrate the feasibility of reconstructing complex subcortical circuits relevant to neuromodulation with dMRI, achieving the most complete such reconstruction with human neuroimaging to date. We validate its anatomical fidelity by mapping topographic organization of cortico-subcortical pathways and replicating connectivity gradients that are consistent with findings from invasive animal studies. Our ultra-high-resolution *ex vivo* dMRI data and pathway annotations, which we make publicly available, open new avenues for training and validating methods for reconstructing these circuits *in vivo* in individual patients. Second, our high-definition, extensible atlas of basal-ganglia-thalamocortical circuits is an invaluable resource for modeling and targeting the effects of neuromodulation in a common coordinate framework. In an initial demonstration of its clinical validity, we show that our atlas captures domain-specific connectivity fingerprints underlying “hotspots” from prior DBS studies.

Previously, we showed that white-matter pathways annotated manually in a small number of subjects with high-quality *in vivo* dMRI data acquired on the Connectome 1.0 MRI scanner could be used to train an algorithm to reconstruct the same pathways automatically from routine-quality dMRI data that could be acquired on conventional MRI scanners^38^. The algorithm used the high-quality training data to learn the anatomical neighborhood of the pathways, which was then leveraged as prior anatomical information that could be applied to dMRI datasets acquired with, *e.g.,* lower b-values or angular resolution. Furthermore, each streamline, rather than each subject, served as a training data point, making it possible to generate millions of training data from a limited number of subjects. Taking this “quality transfer” approach a step further, the high-resolution annotations presented here may be used to train algorithms for automated reconstruction from lower-resolution dMRI that can be acquired *in vivo*. As this approach only assumes that the pathways go through the same anatomical neighborhood in the study subjects as in the training subjects, but not that have the same size, shape, or microstructural features, it may also help bridge the *ex vivo* to *in vivo* divide. This will be a direction for future research enabled by the data that we present here.

Given the challenge of reconstructing brain connectomes in individual patients with conventional human neuroimaging, previous approaches have resorted to constructing normative tract atlases either by aggregating *in vivo* dMRI datasets across large cohorts (*e.g.*, from the Human Connectome Project^39^), or by generating synthetic, idealized streamlines based of the depictions of these tracts in classical anatomic texts^18,19^. Our atlas, constructed from ultra-high resolution *ex vivo* dMRI of human hemispheres, has advantages over both of these approaches.

Normative connectomes built from *in vivo* dMRI data are inherently constrained by limited resolution and SNR. Even the HCP data, which are far superior than typical clinical dMRI scans, are insufficient for reconstructing basal-ganglia-thalamocortical circuits in their full complexity. We demonstrate here that HCP data collected at 1.5 mm resolution fail to reconstruct several small subcortical pathways and exhibit lower accuracy in others, both in terms of their trajectories through the white matter and in terms of their terminations within subcortical structures. The benefit of large *in vivo* datasets such as the HCP is the ability to average across a large N of subjects. While averaging can reduce variance due to noise, validation studies have shown that tractography errors tend to occur at consistent anatomical locations^40,41^, suggesting a systematic bias that cannot be eliminated by averaging alone.

Atlases constructed from synthetic streamlines bypass these issues of conventional dMRI tractography. While such synthesis is done after consulting with classical anatomical texts or with experts on animal tracer studies^18,19^, none of these sources provide a complete, 3D reconstruction of the pathways of interest in the human brain. Thus, synthetic streamlines are, at best, an educated guess fit to a small number of control points.

Our approach leverages the highest-resolution *ex vivo* dMRI data to date from whole human hemispheres, each collected over a week of scanning on the first-of-its-kind Connectome 2.0 MRI scanner to maximize SNR while reaching very high b-values for optimal diffusion contrast. We combine these exquisite datasets with prior knowledge from anatomic studies in human and non-human primates, to annotate both long- and short-range basal-ganglia-thalamocortical pathways, including fascicles with diameters smaller than 2 mm. We present here reconstructions from the first two specimens, and project imaging two additional specimens per year over the next 3 years, which will be used to update our atlas. In addition to the ultra-high-resolution dMRI scans optimized for tractography, which are used to construct this atlas, the same specimens are receiving multi-dimensional (i.e., multi-echo and diffusion time) dMRI scans optimized for microstructural modeling at somewhat lower spatial resolution, presented elsewhere^42^. These can be used to extract estimates of myelination and axon caliber in the same brains, which may be combined with the tractographic reconstructions provided here for advanced modeling of neuromodulation in the future.

Given the challenges in mapping subcortical white matter pathways, previous tractography studies have reconstructed only a subset of the different limbic and motor pathways implicated in neuromodulation across various disorders. The BG circuits, that are critically involved in motor disorders such as PD, DYT, and ET, are composed of both long-range cortico-subcortical projections and subcortical connections. In the present work, we provide extensive reconstruction of both long-and short-range connections within the three BG circuits – the direct, indirect, hyperdirect pathways. To enhance the anatomical and functional specificity of the atlas, we subdivide thalamic, striatal, and subthalamic pathway into subcomponents projecting to distinct regions within the frontal cortex. We pay attention not only to the terminations of the bundles but also to the accuracy of their topographic organization within the IC as they travel from the cortex to their subcortical targets. This is essential for neuromodulation therapies targeting specific sections of the anterior limb of the IC^6,43^.

In addition to the BG pathways, we annotate a substantial portion of the diencephalic and mesencephalic white matter surrounding the BG, including limbic, sensory, and associative pathways. Integrating all major pathways in a single reference framework is essential for providing an updated understanding of the networks modulated in the context of DBS that goes beyond the canonical long-range pathways that currently dominate the field due to the use of lower-resolution dMRI tractography. Alignment of our reconstructed atlas with “hotspots” from prior symptom-mapping studies shows a domain-specific correspondence between patterns of clinical improvement and associated circuits, while highlighting the distributed engagement of motor, sensory, associative, and limbic systems. These results reinforce the anatomical validity of our atlas and highlight its potential utility for both mechanistic investigations and clinically informed interventions.

Beyond DBS applications, this atlas can serve as a foundation for studies of various disease mechanisms and network-level dysfunction, advancing the field by providing more complete and precise models of human brain circuitry. Importantly, the dMRI-based reconstructions presented here are only the first iteration of our atlas, which will be updated as the LINC project collects data with additional modalities, including optical and X-ray microscopy, in the same specimens and in additional specimens. These enhancements will further improve anatomical precision and allow the investigation of inter-individual variability.

## Methods

### MRI data acquisition

The two *ex vivo* human brain hemispheres used in this study were extracted from two different donors at the Massachusetts General Hospital Autopsy Suite and fixed in 10% formalin for at least two months. Demographics are provided in Table 1. Neither donor had known history of neurological disease prior to death. Approximately one week prior to MRI scanning, each hemisphere was transferred into a sealed plastic bag filled with Periodate–lysine–paraformaldehyde (PLP) and checked daily to ensure elimination of most air bubbles that would cause susceptibility artifacts.

**Table 1.**
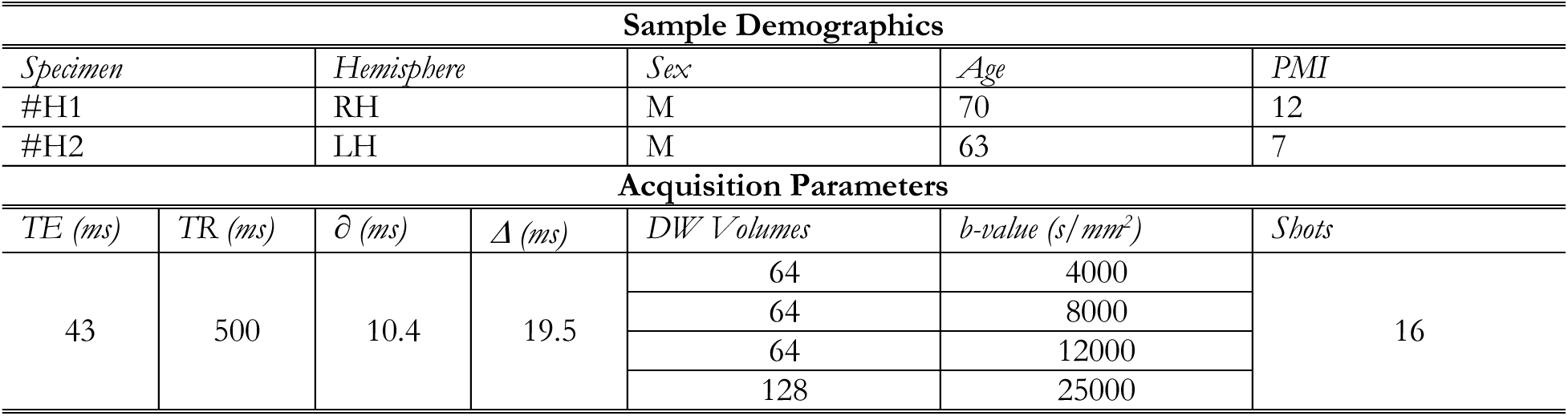
The table reports the demographics for the two samples and the diffusion MRI acquisition parameters. *∂:* gradient pulse width; *1′:* diffusion time; DW: diffusion-weighted; LH: left hemisphere; PMI: post-mortem interval; RH: right hemisphere; TE: echo time; TR: repetition time.

Hemispheres are imaged on the Connectome 2.0, a first-of-its-kind MRI scanner that features the highest gradient strength currently available in a human system (G_max_ = 500 mT/m) [2] using a custom, 64-channel coil array designed for imaging *ex vivo* human brains^44^. The scan consists of 320 diffusion-weighted (DW) volumes with a b_max_=25,000 s/mm^2^ collected at 400 µm isotropic resolution using a multi-shot 3D echo-planar imaging (EPI) sequence^45^ optimized for high-resolution *ex vivo* dMRI^42^ (see Table 1 for details).

### MRI data processing

Data are preprocessed using an in-house pipeline that includes denoising^46^ and correction of image distortions due to B0 inhomogeneities and eddy currents^47,48^ as well as gradient non-linearity. The warp fields estimated for all three types of distortions are combined and applied to each image volume in a one-step resampling process to minimize smoothing effects due to interpolation. Images are then corrected for intensity bias fields and fiber orientation distributions (FODs) are reconstructed using a multi-shell, multi-tissue constrained spherical deconvolution approach (MSMT-CSD) in MRtrix3^49–51^. Prior to the estimation of the tissue-specific response function, a binary map including only white matter and grey matter voxels is obtained using *synthseg* ^52^ to remove CSF voxels. Probabilistic tractography is then propagated from 5 random seed locations in every white-matter voxel^53^ to generate whole-brain tractograms.

Automated segmentation of cortical and subcortical brain structures, as implemented in the FreeSurfer *recon-all* stream^54–57^ (v. 7.5.0) and in SAMSEG^58^ combined with the *Nextbrain* atlas^59,60^, is performed on the spherical average of all volumes collected with b-value = 4000 s/mm^2^, normalized by the average of all volumes collected with b = 0. We replace the parcels of the frontal cortex produced by the aforementioned tools, which are very large, with a refined, publicly available parcellation scheme, comprising the cytoarchitectonic regions defined by Petrides et al. 2012 ^61^ in the space of the *fsaverage* surface-based template^62^. We map that parcellation to the individual surface of each hemisphere using the inverse of the FreeSurfer spherical morph^63^.

### Circuit delineation

We delineate a total of 44 white matter pathways s (Supplementary Fig. 1). Pathways are iteratively generated in a two-step approach (Supplementary Fig. 4). In the first step, the pathways are extracted from the individual whole-hemisphere tractogram using inclusion and exclusion regions of interest (ROIs) in Trackvis. A combination of manually drawn ROIs and ROIs from the automated segmentations are used to guide the extraction of the pathways. Rather than defining the pathways based on the cortical and subcortical ROIs that the fibers terminate in, we use information from prior anatomic tracer studies to identify white matter ROIs that fibers course through. This strategy ensures the accuracy of tractography reconstructions along the entire trajectory of pathways, and not only their terminations. The location of these white matter ROIs is largely guided by the Atlas of the Human Brain^64^. Along with specific basal ganglia pathways, the following surrounding pathways are extracted: the fornix (FX), the stria medullaris (SM), the stria terminalis (ST), the anterior commissure (AC), the mammillo-thalamic tract (MTT), the fasciculus retroflexus (FR), the medial lemniscus (ML), the medial forebrain bundle (MFB), the optic tract (OT), the dentatothalamic tract (DTT). The inclusion ROIs and resulting pathways are meticulously validated by comparison with the anatomical literature, including cadaveric dissection studies in humans^17,65^, axonal tracer studies in non-human primates^13,16,66,67^, histological data from textbooks^64,68^, and supervision by an expert neuroanatomist (SNH). Detailed descriptions of the anatomical definition and delineation of each pathway are included in supplementary section S1.

In the second step, the streamlines from this initial reconstruction of each pathway are converted into a voxel visitation volume that is thresholded to obtain the core of the pathway, and binarized. Tractography is then seeded again from each voxel within this mask; the threshold, number of seeds, maximum tract length, *cutoff*, and *seed_cutoff*, are optimized for each pathway (Supplementary Table 2). Depending on the volume of the initial mask, some tract masks are dilated prior to seeding (Supplementary Table 2).

In addition to white matter pathways, the following ten nuclei are manually delineated in each sample to ensure anatomical accuracy: the internal and external segments of the globus pallidum (GPi and GPe); the subthalamic nucleus (STN); the zona incerta (ZI); the pedunculopontine nucleus (PPN); the substantia nigra (SN); the field of Forel H2; the mammillary bodies (MB); the red nucleus (RN); the habenula (Hb). Manual delineation of these nuclei is performed in FreeView using one or more of the following maps as reference: the mean *low-b* volume (average of all volumes acquired with b-value = 0); the mean normalized *high-b* volume (average of all volumes acquired with b-value = 4000, normalized by the average b = 0 image); the 0^th^-order spherical harmonic coefficient from the MSMT-CSD fit.

### Topographic subdivision of cortical projections

We segment streamlines of the striato-cortical, thalamo-cortical, and subthalamo-cortical projection pathways based on the location of their cortical terminations. We generate frontal ROIs by combining the Petrides^61^ and FreeSurfer cortical parcellations. For some regions, the individual parcels from the Petrides^61^ segmentation are merged to form larger areas. We generate the following subdivisions: dorsomedial prefrontal cortex (dmPFC: area 9M); dorsolateral PFC (dlPFC: areas 946v 46 9L 946d), ventromedial PFC (vmPFC: areas 14, 25), ventrolateral PFC (vlPFC: areas 47, 45 44), orbitofrontal cortex (OFC: areas 14o, 11), OFC/Opro, frontal eye field (FEF, area 8B), premotor (PM, area 6), and motor (M1, area 4). These subdivisions are used to cluster streamlines within the IC, hyperdirect pathway, thalamo-cortical, and striato-cortical projections (supplementary section S1). Streamline endpoints within the STN, thalamus, and putamen are then extracted and converted into volumes. Endpoint volumes in the putamen and thalamus are clustered in FreeSurfer to extract clusters larger than 3 voxels and with streamline density higher than 2. This is repeated for streamlines projecting from each cortical region. We use the putamen clusters as inclusion ROIs to segment the striato-nigral pathway and the thalamic clusters to investigate the relationship between cortico-thalamic projections and subcortical thalamic afferents.

### Atlas creation

We map the white matter pathways and subcortical nuclei from the space of the individual dMRI data to the MNI-2009b-ICBM nonlinear template space (0.5 mm isotropic resolution)^69^, to allow combined analysis with activation volumes from prior DBS studies and to facilitate use of the atlas in future neuroimaging and neuromodulation studies. We use a label-based registration approach to overcome the differences in brain morphology and image contrast between the *ex vivo* dMRI data and the *in vivo* MNI template. We apply the contrast-agnostic segmentation tool SAMSEG^58^, combined with the anatomical designations of the NextBrain atlas to the MNI T1-weighted template. We then use the symmetric normalization (SyN) registration method in ANTs ^70^ to register the segmentation map of the *in vivo* template (masked to include only a single hemisphere) to that of the *ex vivo* hemisphere. The inverse of the resulting non-linear deformation field is used to map the models of the cortico-subcortical pathways and subcortical nuclei onto template space. We warp both the tractography files and their volumetric visitation maps, along with the end ROIs of each pathway. Once in MNI space, we mirror the pathways to the contra-lateral hemisphere and combine them across specimens. We compute the center of gravity for the subcortical nuclei in MNI space and report them in Supplementary Table 3.

### Comparison with publicly available data from the Human Connectome Project

For comparison to our atlas, we annotate the same pathways using publicly available dMRI data from one subject imaged for the MGH-USC HCP ^32^. The dMRI data included four shells (64 directions at b-value = 1,000 s/mm2; 64 directions at b-value = 3,000 s/mm2; 128 directions at b-value = 5,000 s/mm2; 256 directions at b-value = 10,000s/mm2) for a total of 512 diffusion-weighted (DW) volumes and 40 non-DW volumes (b = 0) with 1.5 mm isotropic spatial resolution^32^. Preprocessing is conducted as previously described^38^ and FODs are obtained using the same MSMT-CSD approach in MRtrix3^49–51^. The b=0 s/mm^2^ volume of the HCP subject is aligned to the MNI-2009b-ICBM template using *easyreg* in FreeSurfer^71,72^. The obtained warp is used to map both the voxel visitation maps and end ROIs of each pathway in our atlas to the HCP subject space. Tractography is then performed in subject space using the voxel visitation maps of the atlas pathways as seed regions and the same tract-specific parameters optimized for the LINC data (Supplementary Table 2). For each pathway, we constrain streamlines to go through both end ROIs extracted from the LINC atlas. For the purpose of this comparison, we select a subset of the pathways in our atlas, including both short-and long-range pathways with different levels of anatomical complexity: AL, AS, MTT, ML, OT, DTT, FR, STN-6, STN-4, STN-FEF, ST, SF, and FX. The tractography reconstructions for each pathway are then warped back into MNI-2009b-ICBM template space for visual comparison with those extracted from the LINC data.

### Connectomic analysis of stimulation maps

As further validation of the pathways in our atlas, we use them to identify the circuits associated with therapeutic effects and side effects of DBS. To this end, we curate a set of publicly available probabilistic stimulation maps (PSMs) from previously published studies (Supplementary Table 1). We seek to capture a broad range of DBS indications and stimulation targets across the DBS therapeutic spectrum, and to include maps associated with both clinical benefit and stimulation-induced side effects. In cases where multiple maps are identified for the same indication and anatomical target, a single representative map is chosen based on predefined criteria, including study rigor, patient population (with preference for larger cohorts), spatial specificity of the reported effects, and methodological consistency with contemporary DBS mapping approaches. A total of 18 PSMs are selected by this process, nine of which are associated with clinical improvement and nine with stimulation-induced side-effects (Supplementary Table 1). Maps are resampled into MNI-2009b-ICBM space, thresholded, binarized, and used as inclusion ROIs. For each PSM, we select a threshold that reflects a moderate correlation (or benefit) in relation to clinical outcome (Supplementary Table 1). Annotated streamlines from the atlas intersecting each PSM are then extracted to identify the white matter pathways associated with a given clinical outcome. We calculate the ratio of involvement of each tract by dividing the number of streamlines intersecting the PSM by the total number of streamlines in the tract. To investigate the relationship between symptom domain and associated circuits, we assign each tract in the atlas to either limbic, associative, motor, or sensory domain.

## Data Availability

The LINC atlas is openly available at https://gallery.lincbrain.org/pathways in both individual and MNI152NLIN2009b space. We share the tractography files and their volumetric visitation maps, along with the nuclei at https://dandiarchive.org/dandiset/001278. The MGH-USC HCP are publicly available through the Laboratory of Neuroimaging Image Data Archive (https://ida.loni.usc.edu) and the WU-Minn Connectome Database (https://db.humanconnectome.org).

## Code Availability

Code that generates key results of the study will be available at https://github.com/lincbrain. We used publicly available software tools for the data analysis, including FSL (https://fsl.fmrib.ox.ac.uk/fsl), MRtrix3 (https://www.mrtrix.org), TrackVis (https://trackvis.org), ANTs (https://github.com/antsx/ants), and FreeSurfer (https://github.com/freesurfer/freesurfer).

