## Supplementary Material for "A human subcortical connectome at 400 μm resolution"

Supplementary Information

**Abbreviations**

AC: Anterior Commissure

AL: Ansa Lenticularis

ALIC: Anterior Limb of the Internal Capsule

AS: Ansa Subthalamica

DTT: Dentato-Thalamic Tract

FR: Fasciculus Retroflexus

FX: Fornix

GPe: External segment of the Globus Pallidus

GPi: Internal segment of the Globus Pallidus

H2: Forel’s field H2

Hb: Habenula

Hyp: Hypothalamus

IC: Internal Capsule

LF: Lenticular Fasciculus

MB: Mammillary Bodies

MD: Medial Dorsal nucleus of the thalamus

MFB: Medial Forebrain Bundle

ML: Medial Lemniscus

MTT: Mammillo-Thalamic Tract

OFC: Orbitofrontal Cortex

OT: Optic Tract

Pf: Parafascicular nucleus of the thalamus

PFC: Prefrontal Cortex

Pu: Putamen

PPN: Pedunculo-Pontine Nucleus

RN: Red Nucleus

SF: Subthalamic Fasciculus

SM: Stria Medullaris

SN: Substantia Nigra

ST: Stria Terminalis

STN: Subthalamic Nucleus

Th: Thalamus

VA: Ventral Anterior nucleus of the thalamus

VL: Ventral Lateral nucleus of the thalamus

VLa: Ventral Lateral anterior nucleus of the thalamus

VLp: Ventral Lateral posterior nucleus of the thalamus (also Vim)

VPL: Ventral Posterior Lateral nucleus of the thalamus

VTA: Ventral Tegmental Area

ZI: Zona Incerta

**S1. Manual delineation of pathways**

We report detailed descriptions of region of interest (ROI) placement for the manual delineation of the white matter pathways included in the atlas, along with the anatomical definitions of these pathways. Supplementary Figure 1 shows all the pathways included in the atlas. First, we report the main pathways of the basal-ganglia circuit, then all the surrounding pathways that we included for anatomical completeness. These ROIs represent the protocols used for manual dissections in TrackVis (step 1); the parameters used for pathway refinements (step 2) are reported in Supplementary Table 2. Pathways are listed in alphabetical order.

Basal-ganglia pathways

*Ansa Lenticularis (AL):* This pathway represents the ventral division of the ansa as originally described by von Monakow^1^ (Linsenkernschiinge), also referred to as *ansa lenticularis sensu strictioris* from then after. It is composed of a stratum of thick fibers originating ventrally to the internal segment of the globus pallidus (GPi) that turns dorsally around the medial edge of the internal capsule (IC), runs medial and rostral to the STN, and then mingles with the lenticular fasciculus (LF, described below)^2,3^. Inclusion ROIs were drawn following anatomical landmarks in ^4^: a sagittal inclusion ROI just ventral to the medial pole of the GPi, a coronal inclusion ROI anterior to the STN, and a sagittal inclusion ROI to encompass the ventrolateral thalamus.

*Ansa Subthalamica (AS):* This pathway, described more recently as a separate pathway^5^, connects the ventro-medial GPi to the antero-medial pole of the subthalamic nucleus (STN)^6^, coursing medially, in very close proximity to the fibers of the AL. The STN and GPi were used as inclusion ROIs along with a midway ROI to encompass white matter medial to the AL.

*GPi-Hb:* Fibers leaving the rostral GPi and projecting medio-dorsally towards the thalamic pole to merge with fibers of the stria medullaris (SM) have been identified by virtue of differential staining characteristics in normal and pathological human material ^2^ and in macaques using single axon tracing studies^7^*.* These fibers leave the rostral GPi confined to the periphery of the nucleus and course superior to the pole of thalamus^7^ to merge with the SM^1,8^. Two ROIs were used to delineate the GPi-Hb: an inclusion ROI infero-medial to the GPi (the same used for the AL) and the same inclusion ROI between the anteromedial thalamic nucleus and the lateral ventricle that was used for the SM (described below).

*GPi-Pf:* Fibers leaving the GPi and projecting medio-caudally through the thalamus to reach the parafascicular nucleus (Pf) have been identified in macaques^7^*.* Three inclusion ROIs were used to delineate this pathway: the GPi, a region encompassing the Pf/CM region, and a thalamic waypoint inferior to MD.

*Gpi-PPN (also pallidotegmental tract):* This pathway is formed by fibers that separate from field H and collect dorsal and medial to the STN^2^*.* The fiber group passes ventral to the RN as a band of slightly scattered fascicles, partly mingling with the medial lemniscus (described below), and it terminates almost exclusively in the *nucleus tegmenti pedunculopontinus* (PPN), located inferior, posterior lateral to red nucleus (RN) and dorsal to the SN ^9^. The GPi and PPN were used as inclusion ROIs, along with a third ROI encompassing the white matter ventral to RN.

*Hyperdirect pathway (also Cortico-Subthalamic):* This more recently described pathway was not initially included in the basal ganglia circuit. We delineated the cortical projections to the STN and subdivided them based on their cortical region in streamlines from the dorsomedial prefrontal cortex (dmPFC, area 9M), dorsolateral PFC (dlPFC, areas 946v 46 9L 946d), ventromedial PFC (vmPFC, area 14), ventrolateral PFC (vlPFC, areas 47, 45 44)*,* frontal eye field (FEF, area 8B), premotor (PM, area 6), and motor (M1, area 4) ^1,10,11^. The STN and a ROI drawn on one slice encompassing the anterior limb of the internal capsule (ALIC) were used as inclusion ROIs. Streamlines were then clustered based on their cortical termination regions.

*Lenticular Fasciculus (LF, also fasciculus lenticularis):* This pathway represents the dorsal division of the ansa as originally described by von Monakow^1^. It is composed of fibers that originate caudally to the AL from the medial side of the GPi, traverse the IC, and course between the zona incerta (ZI) and the STN, contributing to the horizontal fiber stratum forming the dorsal capsule of the STN that is commonly labeled field H2 of Forel^1,8^*.*

*Fasciculus Subthalamicus (FS)*: This pathway represents the middle division of the ansa as originally described by von Monakow^1^. Initially described by Willis in 1914 ^12^, these fibers traverse the comb system dorsally extending from the globus pallidus externus (GPe) to the STN, perforating the cerebral peduncle caudal and ventral to the dorsal division (LF). The STN and GPe were used as inclusion ROIs and the GPi was used as an exclusion ROI.

*STN-PPN:* Streamlines connecting STN and PPN were delineated using the STN and PPN as inclusion ROIs, along with a third inclusion ROI encompassing the white matter inferior to the RN.

*STN-SN:* Fibers from the rostral part of the STN descend along the dorsal border of the SN and project ventrally to terminate in the SN ^13^*.* The STN and SN were used as inclusion ROIs and a third inclusion ROI dorsal to the SN was used to isolate streamlines running dorsally.

*Striatonigral pathway:* This pathway, described later than the AL ^14^, comprises fibers projecting from the striatum to the SN. Its course follows the path of the middle division of the AL (*i.e.*, STN-GPe connections) over a considerable distance, and can hardly be distinguished from the latter proximal to the region where both begin to traverse the cerebral peduncle^2^. The SN and putamen were used as inclusion ROIs.

*Striato-cortical pathway:* Striato-cortical connections were subdivided into sensorimotor (areas 4 and 6) and cognitive (dlPFC, vmPFC, vlPFC, dmPFC). The putamen was used an as inclusion ROI, and cortical streamlines going through it were then clustered based on their cortical termination regions: dlPFC (areas 946v 46 9L 946d), vmPFC (area 14), dmPFC (area 9M), vlPFC (areas 47, 45 44), premotor (area 6), and motor (area 4). As the FEF (area 8) mainly projects to caudate, we did not include it ^15^.

*Thalamo-cortical pathway*: The same subdivision used for the hyperdirect pathway was also used for the thalamo-cortical projections, with the addition of 10v, 10ml, OFC, and Opro. The thalamus (as extracted from the *recon-all* automated segmentation) and a ROI drawn on one slice encompassing the ALIC were used as inclusion ROIs. Streamlines were then clustered based on their termination regions.

Surrounding pathways

*Anterior Commissure (AC):* The AC is defined as a fiber bundle running transversely between the anterior part of the bilateral temporal lobes and situated below the fornix medially and the uncinate fascicle laterally^16^. The AC is easily identified in sagittal view; an inclusion ROI was drawn on one sagittal slice in front of the anterior columns of the fornix and a second large inclusion ROI was defined encompassing the white matter of the temporal lobe^17^. The AC was deemed satisfactory after initial manual delineation and did not need refinement through a second seeding step.

*Dentato-Thalamic Tract (DTT):* The DTT courses from the dentate nucleus of the cerebellum through the ipsilateral superior cerebellar peduncle and then partially decussates in the midbrain ^4,18^. Most fibers cross to the contralateral RN and ascend through the posterior subthalamic area to reach the ventral lateral posterior thalamic nucleus^19^. Given that our specimens are hemispheres, we only included ipsilateral streamlines running from the dentate nucleus to the thalamus. The dentate nucleus was extracted from the NextBrain segmentation and used as the first inclusion ROI. A second inclusion ROI was drawn on one coronal slice just dorsolateral to the RN (following landmarks in ^4^) to isolate the streamlines ascending to the VLp. Additional streamlines wrapping around the medial side of the RN and turning anterior to the RN to ascend to the VLp were also observed but not included.

*Fasciculus Retroflexus (FR):* This small pathway courses from the interpeduncular nucleus to the habenula (Hb), medial to the red nucleus. Two inclusion ROIs were used to delineate the FR: the first one was drawn on one axial slice on the medial, anterior, inferior portion of the RN^4^; a second ROI was drawn on one axial slice superior to the RN and inferior to the Hb, at the level of the Pf.

*Fornix (FX):* The FX was defined as streamlines surrounding the thalamus, directly adjacent to the medial half of its superior and posterior surfaces and connecting the hippocampal formation (specifically CA1, CA3, and fimbria) with the anterior thalamic nuclei, the mammillary bodies, the medial septal nucleus, and the basal forebrain ^20^. Four inclusion ROIs were used to delineate the fornix following the anatomical landmarks in^4^: a first inclusion ROI was placed on the axial plane, inferior and posterior to the AC, to outline the column of the fornix; a second and third inclusion ROIs were drawn on one coronal slice anterior to the thalamus and at the level of the Hb respectively to isolate the fornix body; a fourth inclusion ROI was drawn on one axial slice medial to the temporal horn of the lateral ventricle and superior to the posterior hippocampus, at the level of the fimbria.

*Mammillo Thalamic Tract (MTT):* The MTT courses from the mammillary bodies (MB) to the anterior nucleus of the thalamus^4,21^. Its course was clearly visible on the 0^th^-order spherical harmonic image of the MSMT-CSD fit. Two axial inclusion ROIs were used to delineate the MTT: an inferior one just superior to the MB and a superior one posterior and just inferior to the anteromedial thalamic nucleus.

*Medial Forebrain Bundle (MFB):* The classic MFB runs through the lateral hypothalamic area, connects the ventral tegmental area (VTA) to the nucleus accumbens and brainstem, and does not lie within the ALIC but is more ventral in location, remaining inferior to the AC ^4,22^. These unmyelinated fibers are challenging to reconstruct correctly using tractography from conventional dMRI data, but we were able to identify them here. We placed an inclusion ROI at the level of the VTA and two additional coronal ROIs at the level of the MB and optic chiasm, respectively.

*Medial Lemniscus (ML):* The ML ascends through the caudal pons and it gradually slides laterally, so that it lies horizontally in the upper pons, before ascending along the dorsolateral aspect of the RN to reach the VPL nuclei of the thalami. Following descriptions in ^4,23^, we placed a coronal inclusion ROI in the upper pons and two axial ROIs at the posterior inferior and posterior superior level of the RN, respectively. One coronal exclusion ROI was used to discard cerebellar streamlines.

*Optic Tract (OT):* The OT is very clearly visible in coronal view^4^ on the 0^th^-order spherical harmonic image of the MSMT-CSD fit. Three inclusion ROIs were used to delineate the OT: one posterior ROI drawn on one coronal slice over the lateral geniculate nucleus (LGN); one drawn on one coronal slice over the OT at the anterior-most level of the RN; one drawn on one coronal slice at the level where the OT emerges on the midline and starts traversing the brain in a postero-lateral direction. The OT was deemed anatomically accurate and complete after initial manual delineation and was therefore not refined through a second seeding optimization step.

*Stria Medullaris (SM):* The SM courses from the septal area to the habenula, superior to the thalamic dorsomedial nucleus^21^. Following anatomical borders in ^4^, three coronal inclusion ROIs were used to delineate the SM: the most anterior ROI was drawn in the thin space between the anteromedial thalamic nucleus and the lateral ventricle; a second and a third ROI were drawn more posteriorly, medial to the medial dorsal thalamic nucleus.

*Stria Terminalis (ST):* The ST is a major fiber pathway that courses from the amygdala upward and loops around medially in the direction of the hypothalamus. It can be identified in the floor of the lateral ventricle where it accompanies the thalamostriate vein in the groove that separates the thalamus from the caudate nucleus^24^. Three inclusion ROIs were used to delineate the ST: one drawn on one coronal slice at the level of the medial dorsal thalamic nucleus, superior to the reticular thalamic nucleus and medial to the tail of the caudate; one drawn on one axial slice at the level of the lateral pulvinar nucleus between the head and the tail of the caudate; one drawn on one coronal slice posterior and lateral to the LGN following the bend of the temporal horn of the lateral ventricle^4,21^.

Supplementary Figures

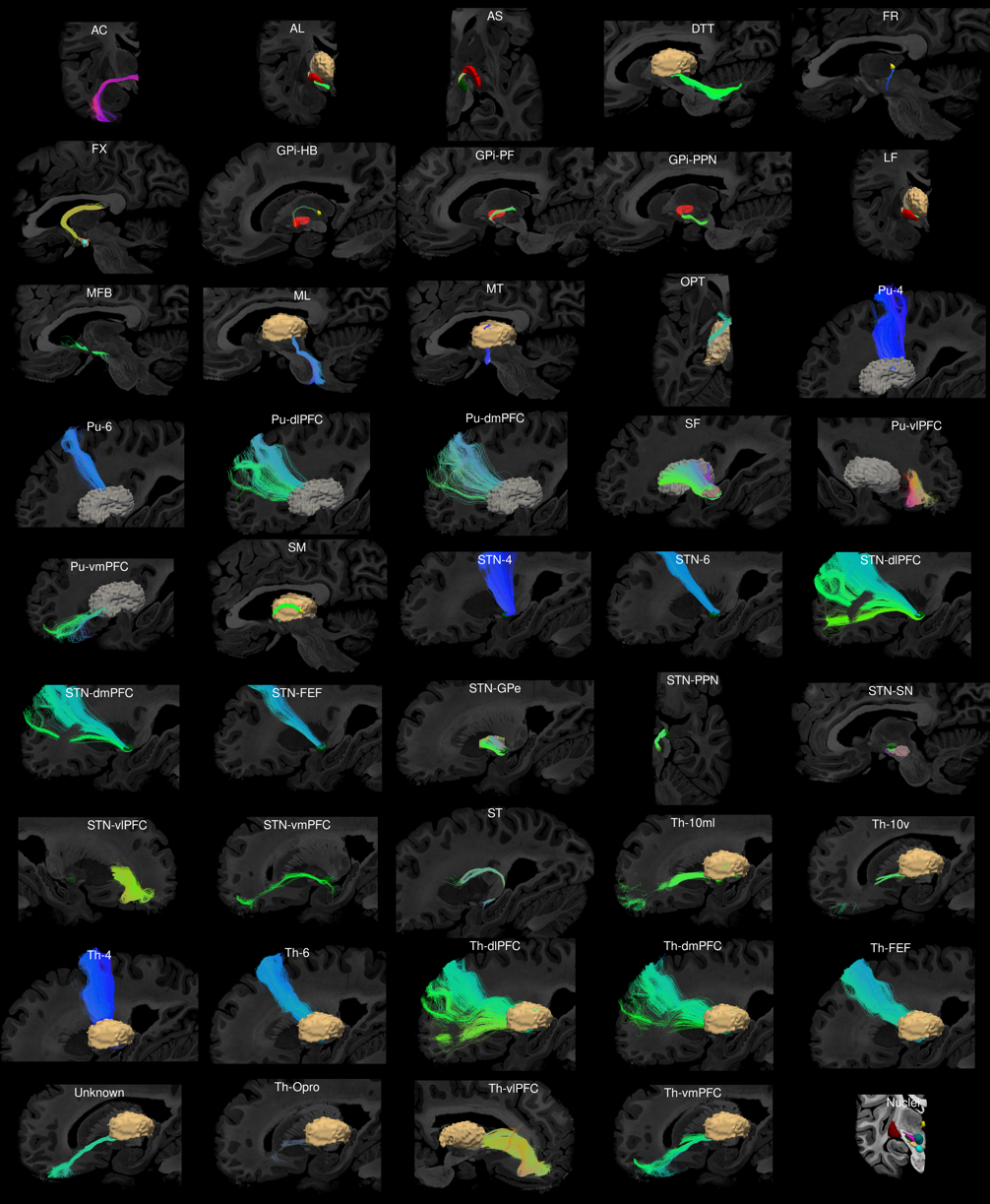

**Supplementary Figure 1. Complete set of atlas pathways.** The figure shows the final 44 delineated pathways in individual space for one specimen (H1). Streamline color encodes directionality (red: medio-lateral; green: anterior-posterior; blue: inferior-superior). 3D reconstructions of related nuclei are also shown for relevant pathways (thalamus in sand, putamen in grey, STN in green, MB in light blue, SN in sand, habenula in yellow, GPi in red, GPe in dark red).

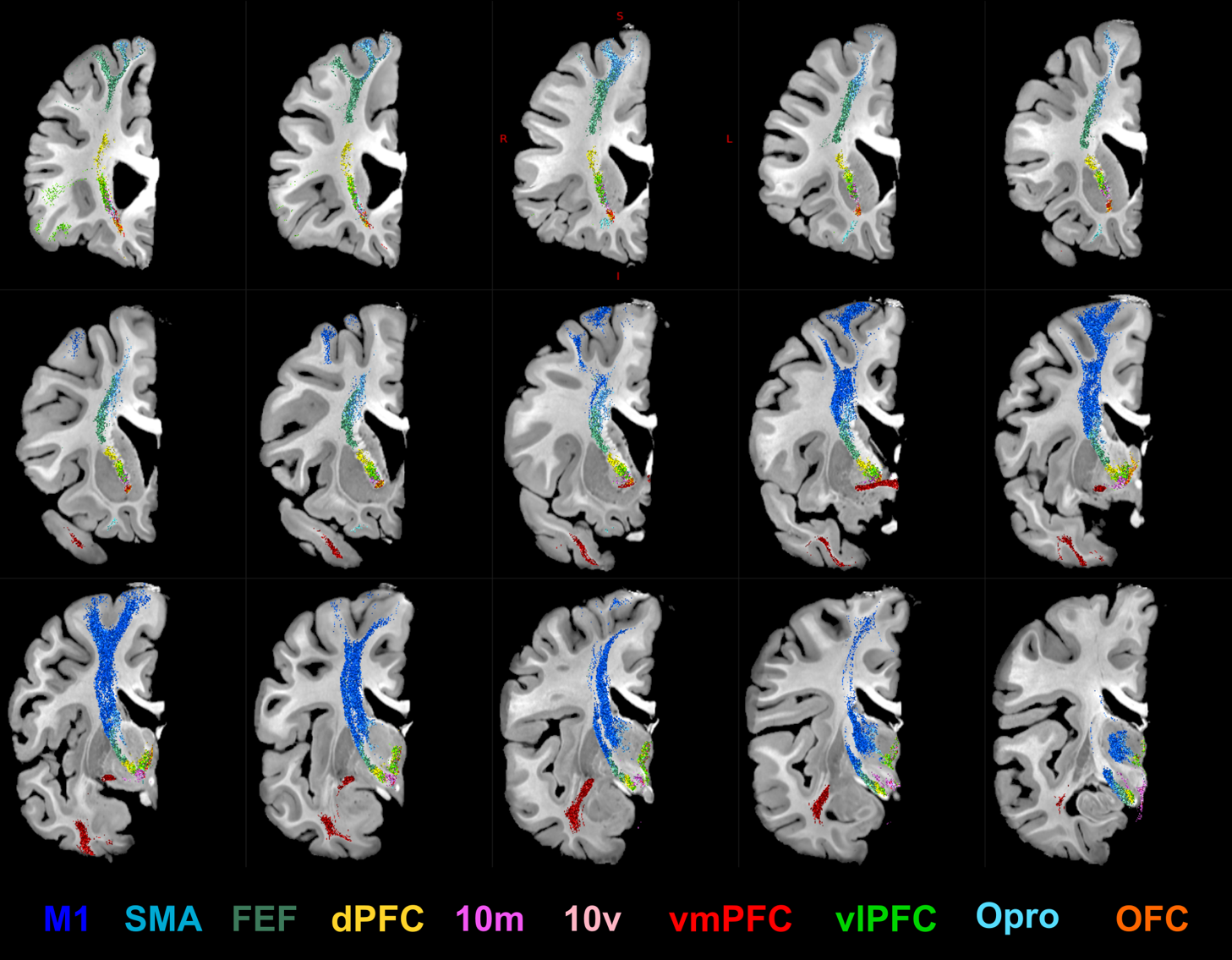

**Supplementary Figure 2. Topographical subdivision of frontal projections for sample H1.** 2D coronal sections showing the location of the different components within the IC and their relationship to the anterior commissure (in dark red). Slices are displayed from more anterior (top left) to more posterior (bottom right) with a interslice interval of 3 mm. Only streamlines intersecting the displayed slice are shown. dlPFC: DorsoLateral PFC; dmPFC: DorsoMedial PFC; FEF: Frontal Eye Field; OFC: OrbitoFrontal Cortex; SMA: Supplementary Motor Area; vlPFC: VentroLateral PFC; vmPFC: VentroMedial PFC.

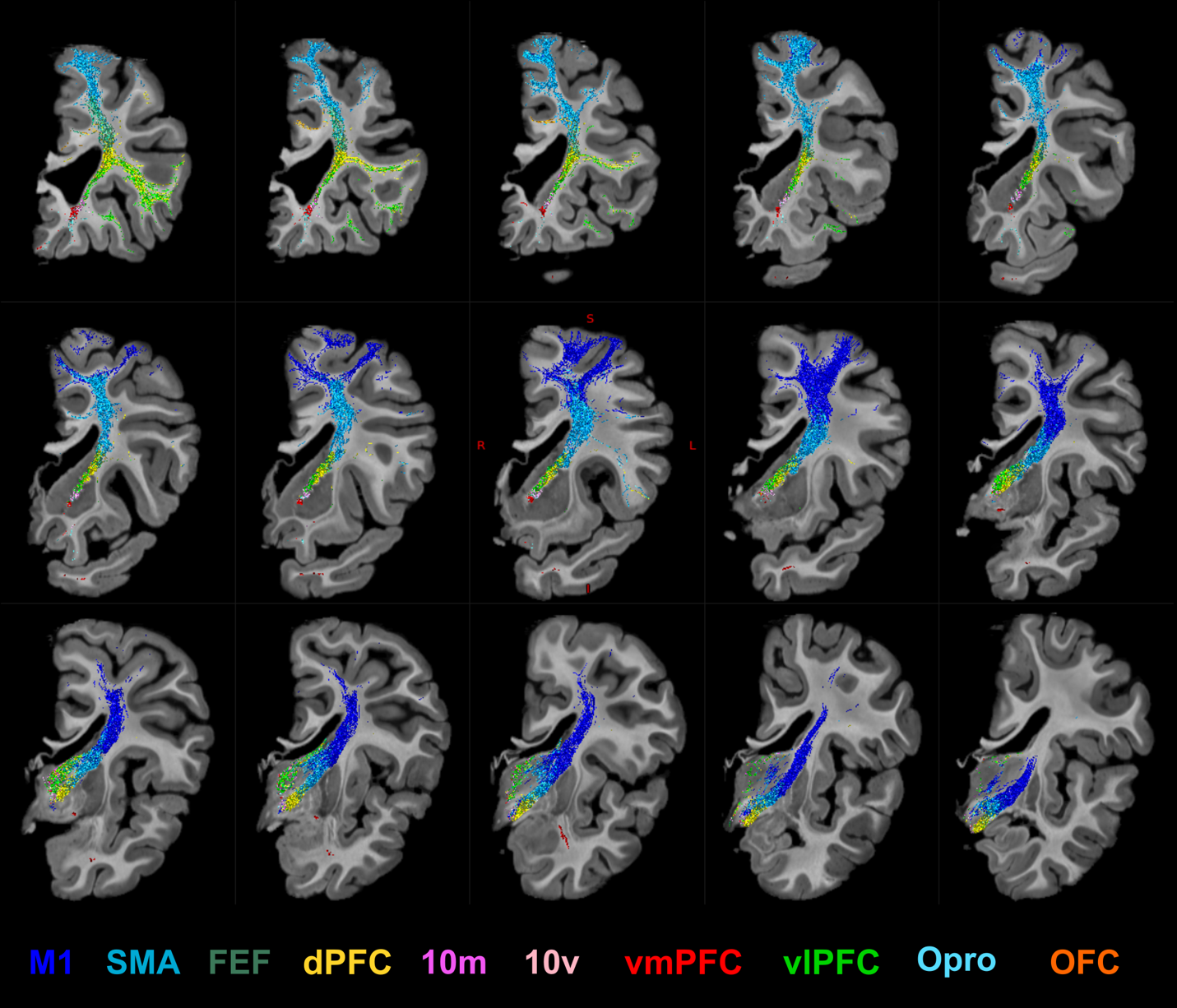

**Supplementary Figure 3**. **Topographical subdivision of frontal projections for sample H2.** 2D coronal sections showing the location of the different components within the IC and their relationship to the anterior commissure (in dark red). Slices are displayed from more anterior (top left) to more posterior (bottom right) with a interslice interval of 3 mm. Only streamlines intersecting the displayed slice are shown. dlPFC: DorsoLateral PFC; dmPFC: DorsoMedial PFC; FEF: Frontal Eye Field; OFC: OrbitoFrontal Cortex; SMA: Supplementary Motor Area; vlPFC: VentroLateral PFC; vmPFC: VentroMedial PFC.

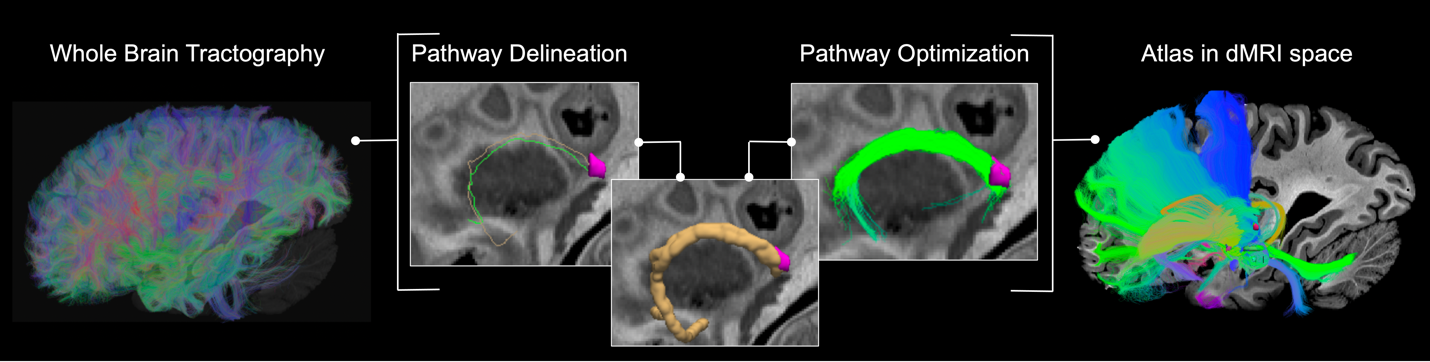

**Supplementary Figure 4**. **Iterative refinement of pathways.** Workflow illustrating the two-step process used to extract and refine white-matter pathways for atlas construction, shown as an example for the stria medullaris. **Left:** Whole-brain dMRI tractography provides the initial set of streamlines. **Middle:** Pathway annotation starts by identifying candidate streamlines from the whole-brain set that course through inclusion ROIs in white matter and terminate in target ROIs in gray matter (*e.g.,* habenula in pink). The selected streamlines are then converted to voxel visitation maps and binarized. Tractography is seeded again from every voxel within this refined mask using optimal tractography parameters that are chosen empirically for each pathway (see supplementary table 1). **Right:** The finalized pathway is integrated into the atlas.

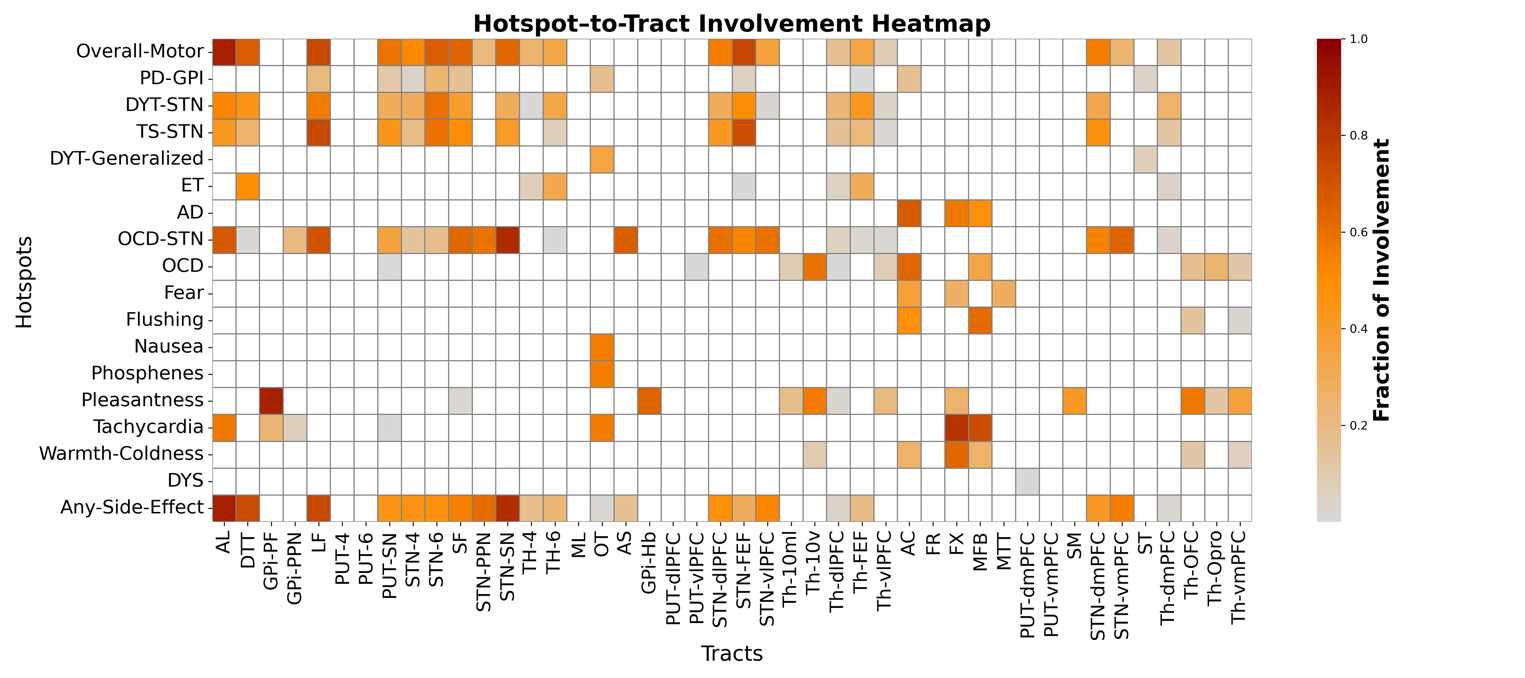

**Supplementary Figure 5. Hotspot-to-tract involvement.** The heatmap reports the fraction of involvement of each tract within the atlas with respect to therapeutic and side-effect hotspot volumes. Details on hotspot maps are provided in Supplementary Table 3.

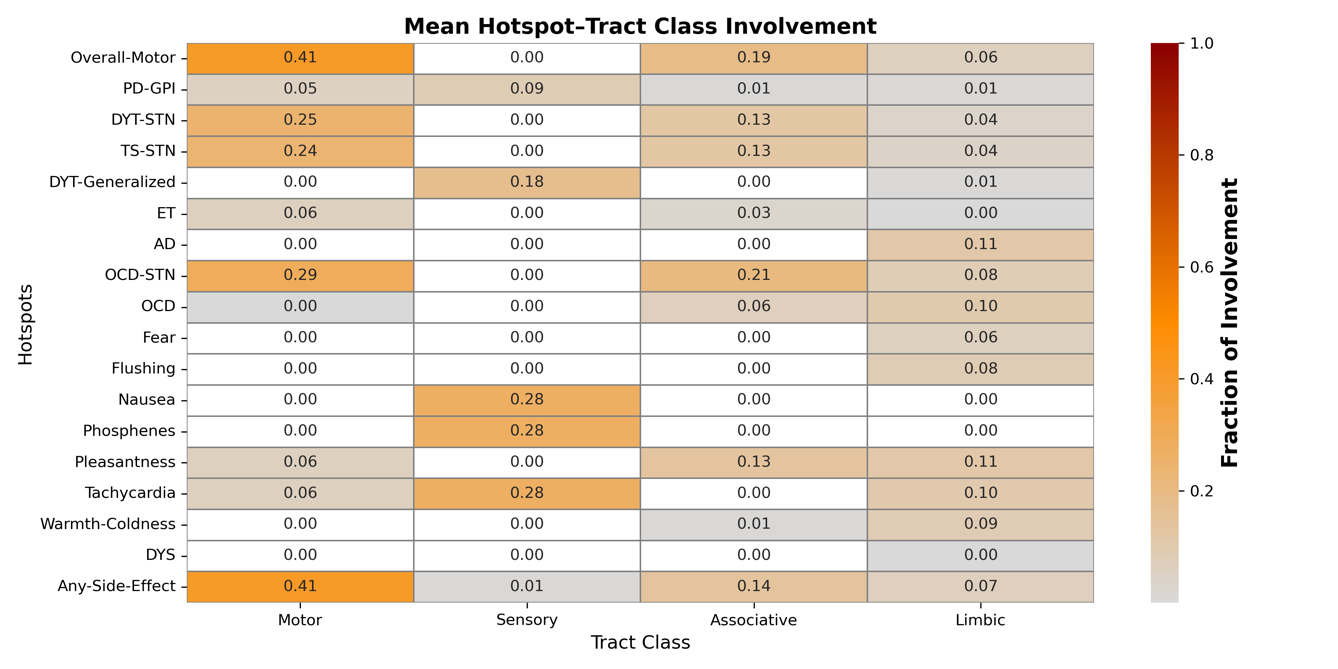

**Supplementary Figure 6. Hotspot-to-Tract Class Involvement.** The heatmap reports the fraction of involvement for each therapeutic and side-effect hotspot averaged over all the pathways within the motor, sensory, associative, or limbic domain. Details on hotspot maps are provided in Supplementary Table 3. Limbic pathways: MFB, STN-vmPFC, STN-dmPFC, Th-vmPFC, Pu-vmPFC, Th-dmPFC, Th-OFC, Th-Opro, Pu-vmPFC, AC, SM, ST, FX, FR, MT; associative pathways: STN-dlPFC, STN-vlPFC, Th-dlPFC, Th-vlPFC, Pu-vlPFC, Pu-dlPFC, Pu-vlPFC, AS, Th-FEF,STN-FEF, GPI-Hb, Th-10ml, Th-10v ; motor pathways: STN-4, STN-6, Pu-4, Pu-6, Th-4, Th-6, Pu-SN, GPi-Pf, STN-PPN, LF, AL, DTT, GPi-PPN, SF; sensory pathways: OT, ML.

Supplementary Tables

| **Type** | **Indication/SE** | **DBS Target** | **n** | **Clinical Outcome** | **Study** | **Stimulation Map** | **Threshold** |
| --- | --- | --- | --- | --- | --- | --- | --- |
| **Improvement** | AD | Fornix |  | ADAS-cog | Rios et al.^25^ | SSM |  |
|  | DYT | GPi | 80 | BFMDRS/TWSTRS | Horn et al.^26^ | SSM |  |
|  | DYT | STN | 70 | BFMDRS | Hollunder et al.^27^ | SSM | 0.2 (moderate correlation) |
|  | ET | Vim | 119 | FTM | Nowacki et al.^28^ | SIM |  |
|  | OCD | STN | 19 | YBOCS | Hollunder et al.^27^ | SSM | 0.2 (moderate correlation) |
|  | OCD | VCVS | 82 | YBOCS | Meyer et al.^29^ | SSM | 0.2 (moderate correlation) |
|  | PD | STN | 21 | UPDRS | Dembek et al.^33^ | MMG | 0.4 (moderate benefit) |
|  | PD | GPi | 28 | UPDRS | Elias et al.^31^ | AMR | 0.2 (moderate hotspot) |
|  | TS | STN | 14 | YGTSS | Hollunder et al.^27^ | SSM | 0.2 (moderate correlation) |
| **SE** | Fear | Hyp | 58 |  | Neudorfer et al.^32^ | VOR | 2 (2x higher odds clinical response) paper’s relevance threshold |
|  | Flushing | Hyp |  |  |  | VOR |  |
|  | Nausea | Hyp |  |  |  | VOR |  |
|  | Phosphenes | Hyp |  |  |  | VOR |  |
|  | Tachycardia | Hyp |  |  |  | VOR |  |
|  | Warmth | Hyp |  |  |  | VOR |  |
|  | Pleasantness | Hyp |  |  |  | VOR |  |
|  | Any SE* | STN | 21 |  | Dembek et al.^33^ | MMG | 0.4 (moderate benefit) |
|  | Dyskinesia | GPi/Gpe | 16 |  | Tsuboi et al. ^34^ | FSL 'cluster' function | Binary.  No threshold |

**Supplementary Table 1. DBS studies included in analysis of stimulation maps.** AD: Alzheimer’s disease ; ADAS-cog 13: Alzheimer’s Disease Assessment Scale—cognitive subscale; AMR: above-mean response; BFMDRS: Burke–Fahn–Marsden Dystonia Rating Scale; DYT: dystonia; FTM: Fahn-Tolosa-Marin tremor rating; Hyp: hypothalamus; MMG: mean map gradient; SE: side effect; SIM: significant improvement map^35^; SSM: sweet spot mapping^26^; TWSTRS: Toronto Western Spasmodic Torticollis Rating Scale; UPDRS: Unified Parkinson Disease Rating Scale; VCVS: ventral capsule ventral striatum; VOR: voxel-wise odds ratio; Y-BOCS: Yale-Brown Obsessive-Compulsive Scale; YGTSS: Yale Global Tic Severity Scale. *Dysarthria, muscle contractions, disturbed vision, dizziness, paresthesia, and mood changes.

| **Tract** | **Max Length (mm)** | **N-seeds** | **FOD Amp.** | **Seed FOD Amp.** | **Threshold** | **Dilate** |
| --- | --- | --- | --- | --- | --- | --- |
| ***AL*** | 30 | 100 | 0.08 | 0.08 | 1 | 1 |
| ***AS*** | 20 | 30 | 0.1 | 0.1 | 1 | 1 |
| ***DTT*** | 100 | 30 | 0.1 | 0.1 | 5 | 1 |
| ***FR*** | 25 | 250 | 0.08 | 0.08 | 10 | 1 |
| ***FX*** | 100 | 50 | 0.08 | 0.08 | 1 | 1 |
| ***GPi-Hb*** | 25 | 200 | 0.1 | 0.1 | 1 | 1 |
| ***GPi-Pf*** | 25 | 150 | 0.08 | 0.08 | 1 | 2 |
| ***GPi-PPN*** | 60 | 100 | 0.08 | 0.08 | 5 | 1 |
| ***LF*** | 25 | 200 | 0.08 | 0.06 | 5 | 2 |
| ***MFB*** | 40 | 250 | 0.06 | 0.06 | 1 | 1 |
| ***ML*** | 45 | 30 | 0.1 | 0.1 | 1 | 1 |
| ***MTT*** | 40 | 50 | 0.08 | 0.08 | 5 | 1 |
| ***NS*** | 50 | 100 | 0.08 | 0.08 | 1 | 1 |
| ***Put-vmPFC*** | 70 | 200 | 0.08 | 0.08 | 1 | 2 |
| ***SM*** | 30 | 50 | 0.08 | 0.08 | 1 | 1 |
| ***ST*** | 90 | 250 | 0.08 | 0.08 | 5 | 1 |
| ***STN-dmPFC*** | 100 | 100 | 0.1 | 0.1 | 1 | 1 |
| ***STN-dlPFC*** | 100 | 100 | 0.1 | 0.1 | 1 | 1 |
| ***STN-FEF*** | 100 | 30 | 0.1 | 0.1 | 1 | 1 |
| ***STN-GPe*** | 20 | 50 | 0.08 | 0.08 | 1 | 2 |
| ***STN-PPN*** | 25 | 250 | 0.08 | 0.06 | 10 | 1 |
| ***STN-SN*** | 20 | 100 | 0.08 | 0.08 | 1 | 2 |
| ***STN-vmPFC*** | 100 | 100 | 0.08 | 0.08 | 1 | 2 |
| ***STN-vlPFC*** | 100 | 30 | 0.1 | 0.1 | 1 | 1 |
| ***STN-4*** | 100 | 100 | 0.08 | 0.08 | 1 | 1 |
| ***Put-vmPFC*** | 70 | 200 | 0.08 | 0.08 | 1 | 2 |
| ***Th-vlPFC*** | 70 | 30 | 0.1 | 0.1 | 1 | 1 |

**Supplementary Table 2. Pathway-specific tractography optimization parameters.** The table reports the optimal tractography parameters that were chosen empirically for each pathway in the second seeding step. For each pathway we set: the maximum streamline length (in mm); the number of random seeds per voxel within the binary mask of the pathway obtained from the first manual delineation step; the fiber orientation distribution (FOD) amplitude threshold used for terminating tracts; the minimum FOD amplitude used for seeding streamlines; the threshold (number of streamlines per voxel) used to binarize the pathway obtained from the first manual delineation step and create the seeding mask for the refinement step; and the number of times the binary mask was dilated.

|  | **STN** | **Hb** | **GPi** | **GPe** | **MB** | **ZI** | **H2** | **PPN** | **RN** | **SN** |
| --- | --- | --- | --- | --- | --- | --- | --- | --- | --- | --- |
| **x** | 9.9 | 1.1 | 17.3 | 19.5 | 0.4 | 10.8 | 6.4 | 4.2 | 3.8 | 10.8 |
| **y** | -12.0 | -23.6 | -5.3 | -1.8 | -9.1 | -12.6 | -8 | -21.6 | -18.6 | -12.6 |
| **z** | -10.1 | -0.7 | -5.2 | -1.2 | -17.7 | -7.2 | -8.1 | -18.8 | -12.7 | -7.2 |

**Supplementary Table 3. Center of gravity of subcortical nuclei.** The table reports the center of gravity for each the subcortical structures included in the atlas. The center of gravity is computed in MNI space and reported in mm.
